# Short Polysialic Acid Counteracts Age-Related Synaptic and Cognitive Deficits

**DOI:** 10.1101/2025.03.13.643031

**Authors:** Loris Frroku, Shaobo Jia, Stepan Aleshin, Srividya Makesh, Katrin Böhm, Aleksandra Alo, Timm Fiebig, Rita Gerardy-Schahn, Markus Fendt, Hauke Thiesler, Alexander Dityatev

## Abstract

Impaired activity of glutamate transporters, elevated concentration of extrasynaptic glutamate and hyperactivity of extrasynaptic GluN2B-containing NMDA receptors are common features in aging and several neurological conditions, including Alzheimer’s disease (AD). Previous studies revealed that polysialic acid (polySia), a glycan predominantly carried by the neural cell adhesion molecule NCAM, inhibits extrasynaptic NMDA receptors and supports synaptic plasticity in healthy adult brains. Moreover, intranasal delivery of polySia with the degree of polymerization 12 (NANA12) rescued synaptic plasticity and cognitive functions in models of tauopathy and amyloidosis associated with AD. Here, we comparatively studied the effects of NANA12 in young (4 months) old (26 months) and very old (29 months) mice. Strikingly, NANA12 promoted cognitive flexibility in attentional set-shifting (ASST) tests and spatial memory in the Barnes maze in very old mice. To capture fine-grained effects undetectable by conventional methods, we introduced a novel trial-wise data analysis approach for evaluating ASST performance. The observed cognitive improvements were not due to changes in the size of hippocampal memory engrams, visualized by c-Fos immunolabeling after reactivation of spatial memory in the probe trial. Five-day treatment with NANA12 did not affect neuronal structure (MAP2 levels), expression of senescence (lipofuscin) or neuroinflammation (microglial Iba1) markers, activation of BDNF receptors (p-TrkB) or expression of endogenous polySia in the hippocampus of very old mice. However, cognitive improvements correlated with the normalized size of CD68^+^ microglial lysosomes and reduced amounts of pre- and postsynaptic proteins at these structures. Thus, our data demonstrate the potential of short polySia to reduce synaptic phagocytosis and restore key cognitive functions attenuated in aging.

## 1. Introduction

According to the United Nations report on global aging, the number of people aged 60 years or older is projected to reach 1.4 billion between 2015 and 2030, while the population aged 80 years or older is expected to triple compared to the number in 2015 (United Nations, 2015) During aging, multiple physiological processes – physical and cognitive – become gradually impaired over time. These progressively impaired processes mostly concern cognitive decline related to memory and other executive functions, such as cognitive flexibility (Peters, 2006; Robbins *et al*., 1998; Smith, 2013). Different brain structures underlying these processes are affected by aging. The hippocampus is a fundamental brain region that enables learning and memory consolidation, as well as spatial memory and other memory-related functions, more generally (Bartsch and Wulff, 2015). It is affected by aging-associated neurobiological alterations. These encompass increased oxidative stress and neuroinflammation, disrupted intracellular signaling, altered gene expression, and reduced neurogenesis and synaptic plasticity, which are believed to contribute to age-related cognitive decline (Bettio *et al*., 2017). In addition to the hippocampus, the orbitofrontal region of the prefrontal cortex is also affected, which plays a crucial role in decision-making, reversal learning, and other higher guiding behavior cognitive functions (Singh *et al*., 2011; Thompson *et al*., 2016).

Age-related cognitive decline has been linked to disruptions in the brain’s excitation-inhibition (E/I) balance, partially through dysregulation of glutamatergic signaling. These disruptions are believed to be associated not only with aging but also contribute to various neural pathologies, including Alzheimer’s disease. Aging may lead to a shift in the E/I balance in favor of uncontrolled excitation and, thus, to neural dysfunction (Ghosh *et al*., 2021). Most excitatory signals in the mammalian central nervous system (CNS) are mediated by glutamate, which plays a crucial role in neural communication, synaptic plasticity, and the regulation of cognitive functions like learning and memory (Iovino *et al*., 2020).

Several aspects of glutamate homeostasis are affected during brain aging resulting in decreased density and function of the excitatory amino acid transporters (EAATs) and leading to reduced glutamate uptake and elevated extrasynaptic glutamate spillover (Segovia *et al*., 2001). Activation of extrasynaptic *N*-methyl-d-aspartate receptor (NMDARs) due to elevated spillover is a contributing factor to excitotoxicity during aging (Magnusson, 2012). It has also been observed that in diseases associated with aging, such as AD, there is a decrease in levels of GluN1, GluN2B, and GluN2A in the synaptic sites and an increase in GluN2B levels at extrasynaptic sites (Escamilla *et al*., 2023). Our previous studies revealed that polysialic acid polymers (polySia) consisting of glycosidically α2,8-linked N-acetylneuraminic acid (NANA) inhibit activation of heterodimeric GluN1/GluN2B-containing NMDARs and heterotrimeric GluN1/GluN2A/GluN2B NMDARs by low micromolar concentrations of glutamate, which are characteristic for the extrasynaptic space (Hammond *et al*., 2006; Kochlamazashvili *et al*., 2010).

Generally, polySia can be considered a collective term encompassing polymers with the degree of polymerization larger than seven (Hayrinen *et al*., 1995; Mindler *et al*., 2021). In the mature hippocampus, polySia associated with the neural cell adhesion molecule (NCAM) is involved in NMDAR-dependent synaptic plasticity (reviewed by (Varbanov and Dityatev, 2017)). Our recent findings show that short polySia with the degree of polymerization of 12 (NANA12), not conjugated to a protein or a peptide, inhibit extrasynaptic GluN2B-containing NMDARs and could rescue impaired synaptic plasticity in murine models of polysialic acid deficiency, tauopathy and amyloidosis, associated with AD (Varbanov *et al*., 2023).

In the present study, we investigated, whether NANA12 can restore cognitive functions impaired as a consequence of aging in wildtype mice. To assess the specific domains particularly involved in cognitive flexibility and spatial memory, we employed well-established behavioral tests for mice that are widely used in experimental neuroscience. Cognitive flexibility was evaluated using the attentional set-shifting task (ASST), while spatial memory, particularly hippocampus-dependent function, was assessed using the Barnes maze test. Interestingly, polySia with degree of polymerization higher than 11 has been reported to bind neurotrophic factors, such as BDNF, which signal via TrkB receptors (Kanato *et al*., 2008). Longer polySia fragments (Schröder *et al*., 2023; Thiesler and Hildebrandt, 2024; Thiesler *et al*., 2022) were found to regulate microglial activity via the polySia-Siglec-axis in mice (Thiesler *et al*., 2021). Here, we verified whether BDNF-mediated TrkB signaling or microglial functions are modulated by NANA12 and may contribute to its cognitive effects in addition to its inhibition of GluN2B-containing receptors. Thus, we assessed the effects of NANA12 in young and very old mice using immunohistochemical analyses to examine potential changes in phosphorylated TrkB and microglial activation. Collectively, our findings provide new insights into the beneficial potential and specificity of NANA12-based interventions for mitigating cognitive aging.

## 2. Materials and methods

### 2.1 Experimental design

During the experiments, experimenters were blinded to the treatment conditions. Mice were divided into different groups across two separate batches, each consisting of 50 % of animals. To ensure balanced experimental conditions, littermates were equally assigned to treatment and control groups. In the first batch, mice were further subdivided into groups of eight, with four tested in the morning and four in the afternoon. Testing was performed in pairs, with each pair receiving equal masses of either NANA12 (treatment condition) or NANA1 (sialic acid or N-acetylneuraminic acid). NANA1 was selected as the control to match NANA12 in terms of charge and composition and as has been reported to have no effects in electrophysiological and behavioral tests in our previous study (Varbanov *et al*., 2023).

During the ASST, mice that failed to complete a specific direct stage of learning were excluded from subsequent stages in a sequential manner: If an animal did not complete the compound discrimination stage (see section 2.5), it was also excluded from the intradimensional shift and extradimensional shift stage testing (six mice treated with NANA1 and five mice treated with NANA12). However, if an animal failed to complete a reversal phase, it was not excluded from subsequent tests.

The Barnes maze test was conducted four days after the completion of the ASST. Out of 47 tested mice, four animals (one old and two very old mice in the control group and one old mouse treated with NANA 12) died during the experimental period and were therefore not tested in the Barnes maze.

During the Barnes maze test, animals were divided into two groups of 12 per batch. There were four subgroups of three animals, each assigned to a different escape hole. Throughout the Barnes maze training of experiment, mice continued to receive the same treatment (NANA1 or NANA12 daily) that was administered during the ASST.

Following the completion of the Barnes maze test, the mice were sacrificed, and their brains were fully extracted for immunohistochemistry. Once all brains were collected, they were analyzed by other experimenters, who were blinded to the age and treatment of the mice.

### 2.2 Animals

Wild type male C57BL/6J mice were used in the experiments, all mice were bred at the animal facility of DZNE Magdeburg. Mice were housed (1-2 per cage) in controlled condition (humidity: 55 ±10%, temperature: 22 ±2°C) with light-on and -off at 6:00 and 18:00 and tested during the light-on period. Twenty-three 3- to 6-month-old males (young group with mean ± SD: 4.8 ± 0.8 months), eleven 25- to 28-month-old males (old group: 25.5 ± 0.9 months) and eleven 29- to 30-month-old males (very old group: 29.1 ± 0.3 months) were used in the experiments. Each age group was randomly split into control and NANA12-treated subgroups, with no significant differences in ages between subgroups. Food and water were provided at *libitum*, only during the ASST experiment, was food restricted (for details see section 2.5). All experiments have complied with the International Guidelines for the Care and Use of Animals for Experimental Procedures (2010/63/EU) with confirmed approval from the local authorities (Landesverwaltungsamt Sachsen-Anhalt, Az. 42502-2-1618 UniMD).

### 2.3 Synthesis of NANA12 and preparation of drug aliquots

NANA12 was generated by enzyme-based catalysis (Keys *et al*., 2014) starting from NANA3 and subsequent purification by anion exchange chromatography applying a DNAPac100 (22 x 250 mm column, ThermoScientific), preceded by a guard column (22 x 50 mm, ThermoScientific). As described (Varbanov *et al*., 2023) elution was detected at 214 nm and isolated NANA12 was desalted applying a size exclusion filtration unit with 2 kDa cut-off (Sartorius, #VS15RH92) and lyophilized The total weight was determined and 200 µg aliquots were prepared by dissolving NANA12 at 2 mg/ml followed by lyophilization. Aliquots were stored at -20 °C until usage. NANA1 aliquots were prepared by the same procedure. Sialic acid polymers were analytically characterized by HPLC-anion exchange chromatography as described before (Keys *et al*., 2014) on a Prominence UFLC-XR system (Shimadzu) using a CarboPac PA-100 column (2 x 250 mm, Dionex) with water and 1M NaCl as mobile phases M1 and M2, respectively, at a flowrate of 0.6 ml/min and column temperature of 50°C. Polymer separation was performed using a -2 curved gradient of 0–30% M2 over 4 min followed by a linear gradient of 30–84% M2 over 33 min and detection at 214 nm. (Figure 1). NANA1 was obtained from Molekula Group (#596039, batch #205711), disodium cytidine 5’-triphosphate (CTP) was from Sigma/Merck (#C1506-1G, batch #102515151) and sialic acid trimer (NANA3, #00641-52, lot #Z4N7809) and sialic acid teramer (NANA4, #00642-42, lot #Z5B8604) were from Nacalai Tesque, Japan.

**Figure 1.**
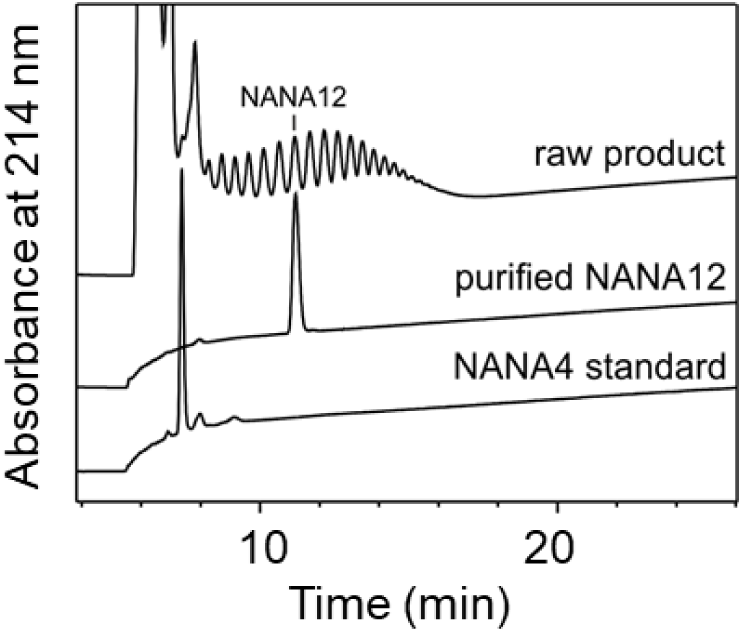
Anion exchange chromatography of N-acetylneuraminic acid (NANA) polymers. The signals correspond to a raw product of enzyme-based catalysis before preparative anion exchange chromatography, purified NANA12 after aliquoting and lyophilization and commercially available NANA4 standard. Detection is based on absorption at 214 nm.

### 2.4 Drugs

Mice received NANA12 or, as a control, NANA1 (free sialic acid) in saline solution. Drugs were delivered intranasally (Varbanov *et al*., 2023) via four consecutive 4 µl injections (1 mg/ml) in each nostril of the mouse (32 µl per mouse in total) two hours before the following experimental setups: compound discrimination, intradimensional shift, extradimensional shift during ASST and before the first training session of every training day in the Barnes maze experiment.

### 2.5 Attentional set shifting task (ASST)

Mice were subjected to a well-established paradigm of ASST to study cognitive flexibility. It is sensitive to prefrontal cortex-mediated cognitive deficits (Heisler *et al*., 2015). In this paradigm, animals learn which stimulus dimension is relevant by trial and error, receiving positive feedbackthrough a reward (Brown and Tait, 2010). An attentional set is formed when a subject learns that a set of rules can be applied to complex stimuli in order to differentiate relevant from irrelevant cues. Animals learn to pay attention to a relevant stimulus dimension and ignore another one during a series of discrimination tasks: compound discrimination, intradimensional shift, and extradimensional shift. Animals also perform the reversal learning tasks at all these stages (Brown and Tait, 2010).

#### Set up and materials

The ASST custom-made box (41 cm x 22 cm x 24 cm, University of Magdeburg, Germany) was slightly modified in comparison to a setup described elsewhere (Seifried *et al*., 2023) to include an additional waiting compartment separated from two choice compartments by opaque plastic walls with a sliding door. The door could be opened by the experimenter to allow an animal to pass between the compartments but to prevent observing the content of the respective choice compartments before the start of a trial, as shown in the Figure 2A. The two-choice compartments were separated from each other by a transparent plastic wall. In each of the compartments, a bowl was placed (5 cm diameter, 2.5 cm height). The bowl in the waiting compartment was filled with water, while the bowls in the choice compartments were filled with 4 distinct media with approximately the same size, and 4 diverse odors were applied during the different stages of the ASST, media 1 (M1) was brown clay spheres; M2 was green wood spheres; M3 was fluff red spheres; M4 was aluminum spheres. Four odorants were used in a 1:20 dilution in paraffin oil, such as O1: valeric acid; O2: S-(+)-carvone; O3: citral; O4: R-(+)-carvone. 40 µl of the odors were applied on a filter paper every 2 h to refresh it. The filter paper was attached to the front of the bowl.

**Figure 2.**
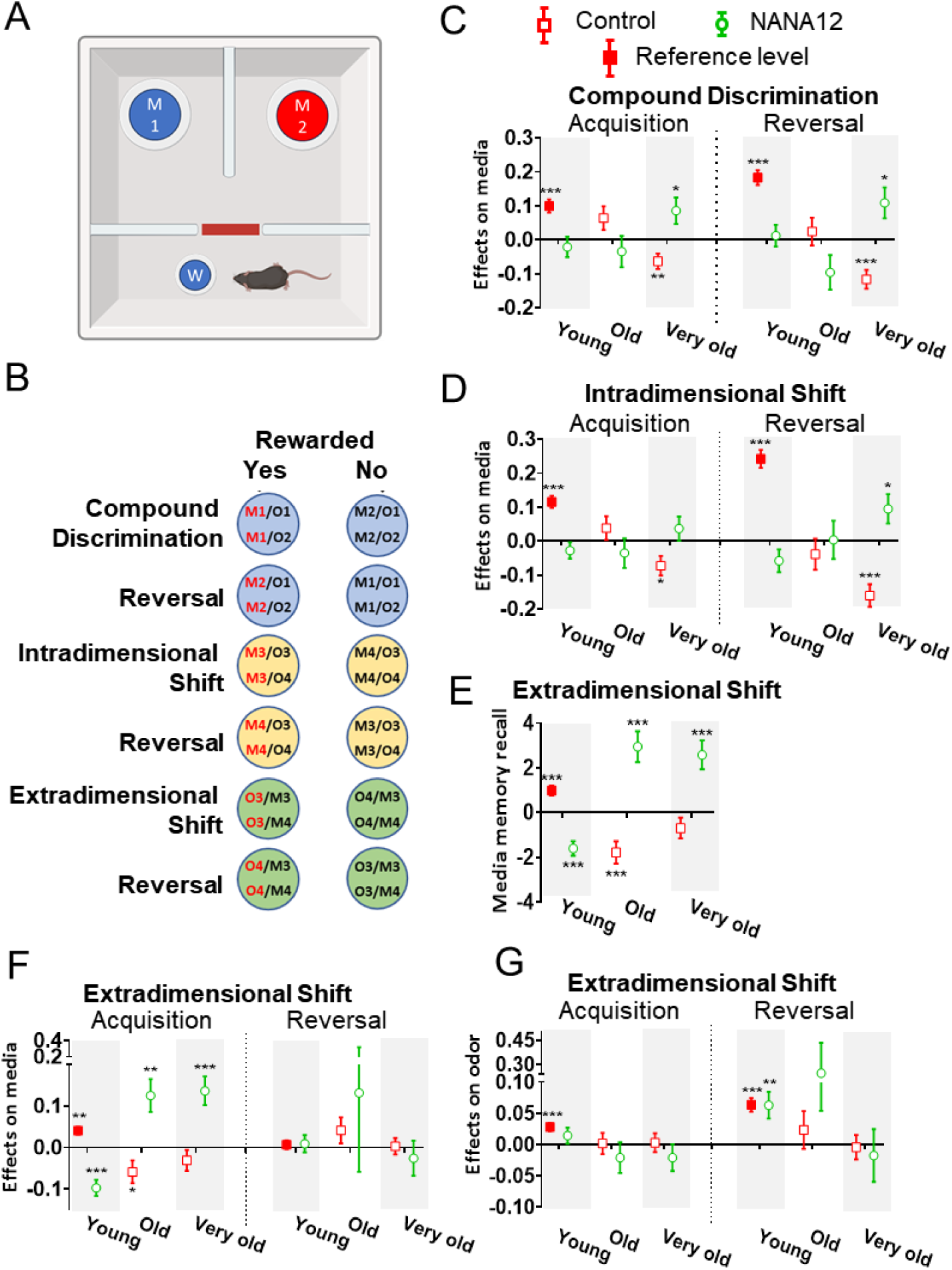
Age- and drug-related differences between groups in ASST. **(A)** Apparatus for ASST. **(B)** Scheme of experiment. **(C)** Compound discrimination stage, showing learning and reversal learning rates for medium 1 and 2, correspondingly. **(D)** Intradimensional shift stage with novel exemplars medium 3 and 4 learning rates. **(E)** Medium memory recall at extradimensional shift onset, indicating how strongly mice expect medium 4 (vs. 3) to be rewarded. **(F)** Extradimensional shift stage with medium 3/4 learning rates. **(G)** Extradimensional shift stage, learning and reversal learning rates of odors 3 and 4. Error bars denote standard error. Significance thresholds are indicated by asterisks (\**P* < 0.05, ** *P* < 0.01, *** *P* < 0.001).

#### Handling and food restrictions

From day 1 to day 3 animals went through handling habituation to the experimenter. Animal weight was measured every training day and at the end of the third day, an average of the weights was calculated and considered as the mouse basal weight. From day 4, food supply was restricted, and only approximately 2.5 g/day of food was given to single-housed mice to maintain 90-95% of their basal weight. Food was replenished daily following behavioral testing.

Starting on day 4, mice were acclimatized to the digging bowl and the associated reward. A digging bowl was placed inside the mouse cage, with a reward (choco rice from pellet, Nordgetreide GmbH, Luebeck, Germany) placed at the bottom of the bowl and covered with bedding material. Additional rewards were scattered on top of the bedding in the bowl and around the cage to facilitate the animals’ familiarization with it.

#### Buildup stages

The buildup stages were designed with the intention to habituate the animals to the ASST setup, as well as to the various media and odors they would encounter during the actual test. During these stages, all possible combinations of odors and digging media were prepared and systematically introduced.

Buildup Stage 1: In this stage, each mouse was tested for 1h. Mice were placed in the waiting compartment of the ASST setup, and digging bowls were randomly placed in the choice compartments. Both bowls were rewarded, with the reward located at the bottom of the digging media. Once the sliding door was opened, the animal was allowed to freely explore the compartments. A trial was considered concluded when the mouse actively dug in both bowls and retrieved the rewards and then moved back into the waiting compartment. For each new trial, a different odor/media combination was used. Digging behavior was defined as the active displacement of the media using either the paws or nose to access the reward.

Buildup Stage 2: After a 48h break, this stage was conducted over two consecutive days. Unlike Buildup Stage 1, the trials involved bowls that were either rewarded or non-rewarded, with the condition randomly determined for each trial, with a limit of three maximum consecutive rewarded or not rewarded trials in a row. As in the previous stage, the trial ended once the mouse had dug in both bowls. Each testing session lasted one hour.

#### Assistance during buildup phases

During both buildup stages, assistance was provided to the animals if, during the first trial, they were unable to locate the reward or were not motivated to dig. In these cases, a small reward was placed on top of the digging media to encourage exploration and digging behavior. These buildup stages ensured that the animals became adapted to the ASST setup, the digging task, and the various experimental contingencies.

#### Compound discrimination and reversal

During the compound discrimination stage, animals were presented with stimuli consisting of two choices of the relevant dimension (media) and two of the irrelevant dimensions (odors). To avoid an animal to possibly smelling the reward, a light dusting from the same choco rice pellets used to reward the mice was applied on the top and the bottom of the bowls. The relevant dimensions included M1 (rewarded) and M2 (non-rewarded), while the irrelevant dimension included O1 and O2. These stimuli were paired in the following combinations: M1/O1, M1/O2, M2/O1, and M2/O2. This stage lasted 1 hour, during which animals were tasked with learning through trial and error. Only the media dimension was associated with a reward, while the odors were irrelevant. Within the media dimension, only M1 was rewarded. If an animal dug correctly in the rewarded bowl (M1) in 8 out of 10 consecutive trials, the compound discrimination stage was concluded, and the animal advanced to the compound discrimination reversal stage with the same 1h limit to learn the task. During the compound discrimination reversal stage, the reward association was reversed: M2 became the rewarded media, while M1 was no longer rewarded. These stages tested the animal’s ability to learn and adapt to the relevant stimulus dimension and to adjust their behavior when reward contingencies changed. Between tests of different mice, the ASST setups were cleaned with 70% ethanol to avoid any odor to distract the following mouse.

#### Intradimensional shift and reversal

During the intradimensional shift and its reversal, the training rules followed those outlined in the compound discrimination section. In the intradimensional shift stage, the mouse was presented with novel stimuli providing two choices in the reward-relevant dimension (media): M3 (rewarded) and M4 (non-rewarded) and two choices in the irrelevant dimension (odors): O3 and O4. These stimuli were paired in the following combinations: M3/O3, M3/O4, M4/O3, and M4/O4. While the specific media and odors differed from those in the compound discrimination stage, the relevant dimension (media) was maintained. During the intra-dimensional shift, M3 was the rewarded media. During the followed reversal stage, the reward contingency was reversed: M4 became the rewarded media, and M3 was no longer rewarded.

#### Extradimensional shift and reversal

During the extradimensional shift for the first time, the previously relevant dimension (media) became irrelevant, i.e. the correct choice was a cue previously associated with the reward-irrelevant dimension (odors). In the extradimensional shift stage, a mouse was presented with the same media and odor combinations used in the intradimensional shift acquisition and reversal stages: M3/O3, M3/O4, M4/O3, and M4/O4. However, the reward association shifted to the odor dimension. Specifically, O3 was rewarded during the extradimensional shift stage. During the extradimensional shift reversal stage, the reward contingency reversed within the odor dimension: O4 became the rewarded cue, and O3 was no longer rewarded. The timing and stopping criteria for both extradimensional shift acquisition and reversal stages were consistent with those described for the compound discrimination. These stages tested the animals’ ability to shift their attention to a completely new relevant dimension while adapting to reversed reward contingencies.

### 2.6 Statistical evaluation of ASST data

We fitted two main generalized linear mixed models (GLMM) to our binary choice data (left vs. right bowl), each capturing different stages of the ASST. We used a binomial distribution with a logit link (via `fitglme` function in MATLAB 2024a), allowing random intercepts by mouse (*mouseID*) to account for individual biases in left/right and stimulus preferences.

Each trial was represented by a single row in the dataset, indicating whether the mouse chose the left bowl (*isLeft* = 1) or the right bowl (*isLeft* = 0). We included two trial counters: *directTrialCount*, which increased only during the direct (initial) stage of each stage, and *reverseTrialCount*, which increased only during the reversal stage. This allowed the model to estimate separate slopes for direct versus reversal learning.

To encode stimulus-reward contingencies, we introduced two key predictors: *directMedLeft* and *directOdorLeft*, each taking values of +1 if the left bowl’s content (medium or odor) was the rewarded option in the direct stage, and −1 if it was the option rewarded in the reversal stage.

We also included three binary factors—*isOld*, *isVeryOld*, and *isDrug*—to reflect whether the mouse was in the old or very old group, and received NANA12 treatment, respectively. By incorporating *(isOld + isVeryOld)×isDrug* in the fixed-effects structure, we could assess main effects attributed to the dimensions age or treatment and any interactions.

Additionally, we accounted for individual differences in baseline stimulus preference by introducing two indicators—*O1/3* and *M1/3*—which did not change over trials but instead captured each mouse’s initial preference for the stimuli used in the current phase. These indicators were treated as random intercept-like offsets: A mouse might exhibit a stable preference for a particular stimulus from the outset of the direct stage. Thus, to reflect these individual-level differences, we allowed each mouse (*mouseID*) to have a random intercept (capturing left/right biases) as well as random intercept-like parameters for odors (*O1/3*) and media (*M1/3*). These indicators were considered equal to 1, when in the left bowl was corresponding stimuli, and zero, when the corresponding stimulus was in the right bowl.

Taken together, for compound discrimination and intradimensional shift stages we used the following formula:

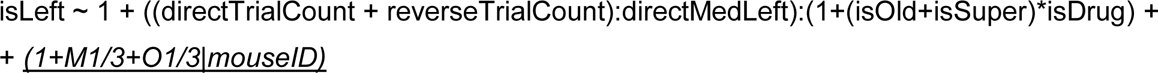

where -1 excluded a global intercept (thus, in the absence of any predictors, the probability of choosing left was assumed to be 0.5 overall). In these stages, the relevant dimension (medium) did not change, so mice were presumed not to carry a strong group-level prior from a different dimension; any baseline preference was absorbed by the random intercepts and by the stimulus-specific intercepts *O1/3* and *M1/3*.

We found that this model explained a substantial portion of the variance in choice: For compound discrimination, the AUC was 0.85 and R^2^ = 0.36; for intradimensional shift, AUC was 0.84 and R^2^ = 0.34, (p ≪ 10^−6^ for both models).

The extradimensional shift stage introduced a change in the relevant dimension from medium to odor. By this point, mice had learned to discriminate based on medium in intradimensional shift, so they might carry a systematic bias favoring the formerly correct medium into the early trials of extradimensional shift rather than starting from a 0.5 baseline. To capture that effect, we specified a more comprehensive fixed-effects block in the extradimensional shift formula:

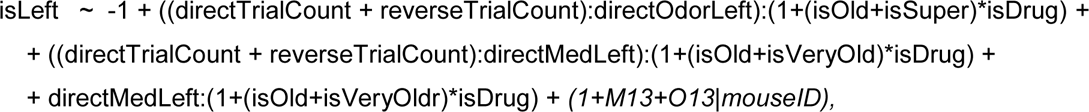

allowing *directMedLeft* to influence extradimensional shift choices from the outset. Interactions between *directMedLeft* and the trial counters (*directTrialCount*, *reverseTrialCount*) enabled the model to reveal whether the mice’s prior reliance on “correct” media from the intradimensional shift stage decayed or persisted over extradimensional shift trials.

The extradimensional shift model continued to perform well (AUC ≈ 0.81, R^2^=0.29, p≪10^−6^), though slightly lower compared with compound discrimination/intradimensional shift, suggesting that shifting to a new dimension posed a greater challenge.

For all models, we employed `CovariancePattern = ‘Full’` to allow correlated random terms, `CheckHessian = True` to ensure robust convergence checks, and `DispersionFlag = True` to handle potential overdispersion in the binomial data.

The statistical significance of fixed effects was determined by examining fitglme output. The model coefficient estimates, confidence intervals, and further discussion of how age (*isOld*, *isVeryOld*) and drug treatment (*isDrug*) influenced behavioral performance are given in the Results section (Figure 2).

To complement these results, we provide additional supplementary visualizations. Figure S1 illustrates individual learning trajectories alongside model predictions, offering insight into variability in performance. Figure S2 presents the estimated learning rates for each group, along with post-hoc statistical comparisons. While Figure 2 highlights how treatment and age influence learning dynamics, Figure S2 provides a broader perspective on overall group performance by displaying absolute learning rate estimates.

### 2.7. Barnes maze

### The Barnes maze setup

The setup of the Barnes maze consists of a custom-made plastic disk with a diameter of 92 cm, featuring 20 holes, each with a diameter of 5 cm, distributed around the platform (University of Magdeburg, Germany). These holes are positioned 2.5 cm from the outer edge and allow connection to an escape box located underneath the platform, which can be placed under the hole chosen by the experimenter (Figure 3B).

**Figure 3.**
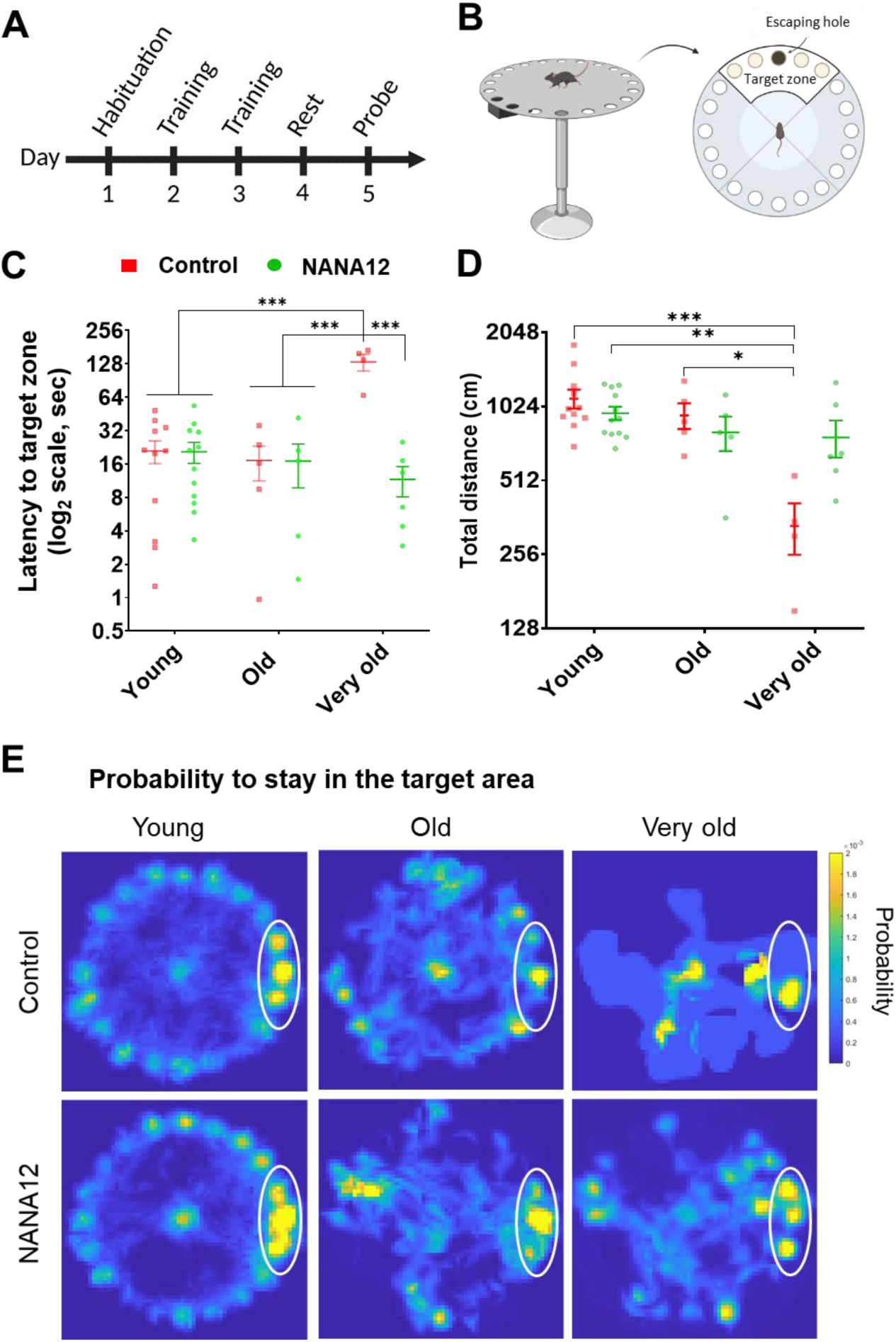
NANA12 improves spatial learning of very old mice in Barnes maze. (A) Simplified diagram representing the timeline of the Barnes maze experiment. (B) Schematic representation of the apparatus used for the Barnes maze test, showing the target zone and escaping hole. (C) NANA12 decreases the latency to reach the target zone in very old mice (n=6) compared to very old control mice (n=4). (D) NANA12 increases the distance moved by very old mice compared to very old controls, resetting it to the levels of old and young mice. Data are presented as mean ± SEM on a logarithmic scale. Data were obtained using a two-way repeated measures ANOVA, followed by Tukey’s post hoc test (*p<0.05, **p<0.01, ***p<0.001). (E) Old and very old mice treated with NANA12 preferentially stayed in the area near to the escape hole during the probe trial.

The setup is placed in a chamber, and various visual cues depicting colored geometric shapes are positioned around it. There is no access to natural light from the outside to prevent variations in light intensity within the room. Instead, eight lamps provide a controlled environment with an illumination level of 912 lux, as determined by a luminometer. During the Barnes maze test, the animals were recorded using the CyberLink YouCam software, version 5.0. Subsequently, the videos were analyzed using the EthoVision 11.0 XT software by Noldus (Figures 3C and 2D) and the DeepLabCut software (Figure 3E).

### Habituation and introduction to the task

The mice faced the Barnes maze in three stages: habituation, training, and probe trail as described elsewhere (Attar *et al*., 2013) with minor changes. On day 1, mice were habituated by placing in the center of Barnes maze inside a transparent plastic cylinder to allow the animal to see the surroundings for 30 s. Then by gently moving the plastic cylinder, mice were guided to the escaping hole in the target zone that leads to the escaping box (Figure 3B). Then, mice were given 1 min to enter the escaping box independently. If an animal did not enter the escape box by that time, it was gently guided in the cylinder to the escape hole and forced to enter the hole. The mice were left in the escaping box for 1 min before returning them in their cages.

### Training

The training stage lasted for two days. Each mouse performed 3 trials per training day with a 20-min inter-trial interval. Mice were placed inside an opaque cylinder in the middle of the Barnes maze for 10 s, to avoid animal using visual cues around the room and to face a random direction when cylinder was lifted, and the trial began. After the cylinder was removed the mice had 3 min to explore the maze. If a mouse found the target hole and entered the escape box at that given time, the trial was considered over and the animal was allowed to stay inside the escape box for 1 min, before being returned to the home cage. If a mouse did not enter the escape box at that time, it was guided with the transparent cylinder to the escaping hole and let to enter the escape box independently and let inside for 1 min. Between trials the platform and the escape box were cleaned with 70% ethanol, to avoid any odor to be distracting for the following mouse. The probe trial was performed 48 hours after the last training day.

### Probe trial

During the probe trial, the escape box was removed, and mice were placed inside an opaque cylinder in the middle of the Barnes maze for 10 sto avoid animal using visual cues around the room and to face a random direction when the cylinder was lifted, and the trial began. Like on the training day, the animal was allowed to explore the platform for 3 min at the end of which the mouse was returned directly to the cage. The probe stage was video-recorded and measures of the latency to reach the target zone and time spent there were analyzed.

### 2.8 Brain sample collection

Ninety min following the completion of the final stage of the Barnes maze (probe trial), mice were anesthetized with a combination of ketamine (100 mg/kg), xylazine (20 mg/kg), and acepromazine (3 mg/kg). Intracardiac perfusion was then performed for 2 min with phosphate-buffered saline (PBS, pH 7.4; Thermo Fisher), followed by 10 minutes with 4% formaldehyde solution (Roti-Hotfix 4%, pH 7; Carl Roth GmbH + Co. KG). After perfusion, animals were decapitated, and whole brains were extracted and immersed in 5 mL of 4% formaldehyde overnight before proceeding to further analysis.

### 2.9 Brain section preparation

Brain sections were prepared similarly to our previous studies (Ferrer-Ferrer *et al*., 2023). After postfixation with 4% PFA overnight at 4 °C, brains were incubated in 30% sucrose/PBS solution for 2 days to cryoprotect the tissue. Finally, the brains were frozen in 100% 2-methylbutane at -50 °C and then stored at -80 °C. For cryosectioning, brains were embedded in the tissue freezing media from Leica (#14020108926) and cryo-sectioned into 40-μm-thick coronal sections (Leica CM 1950, Leica). Floating sections were kept in cryoprotective solution (1 part of ethylene glycol, 1 part of glycerin, and 2 parts of PBS, pH 7.4).

### 2.10 Immunostaining

For general immunostaining, brain sections were washed 3 times in 1X phosphate buffer (PB) pH of 7.2 for 10 min with gentle shaking; then incubated in blocking and permeabilization solution (BP solution: 5% (v/v) normal goat serum, 0.3% (v/v) Triton X-100, 0.1% (w/v) glycine in 1x PB) for 1 hour, followed by primary antibody incubation in BP solution (350 μl per well in 48-well-plate) at 4°C for 2 days (Table S1). Later, all sections were washed3 times in 1x PB and incubated with secondary antibody (in BP solution) at room temperature for 3 hours, then washed twice in 1x PB. To stain nuclei, sections were incubated with DAPI at 1:400 for 10 minutes. Subsequently, sections were briefly washed in ddH_2_O to remove salts and mounted on Superfrost® plus microscope slides (Epredia, #J1800AMNZ) with Fluoromount (Sigma, #F4680).

### 2.11 Image capturing

Images were acquired using an inverted microscope (CellDiscoverer 7 with LSM-900, Carl Zeiss) or an upright microscope (LSM 700, Carl Zeiss), depending on experimental purpose and availability. When using CellDiscoverer 7, widefield mode was selected to generate a panoramic navigation map, and then tile-scanned images with higher resolution under widefield mode or confocal mode were acquired for brain region-specific marker intensity analysis or cell counting. When using LSM-700, images for cell counting were acquired with low magnification objective lens; Z-stack image sets for microglia volume and synaptic pruning analysis were acquired with a higher magnification objective lens.

The objective lenses used on the CellDiscoverer 7 were a Plan-Apochromat 5x/0.35 NA air objective, a Plan-Apochromat 20x/0.95 NA air objective and an optovar with 0.5x/1x/2x tube-lens; the objectives used on the LSM-700 confocal microscope were an EC Plan-Neofluar 10x/0.3 NA air objective and a Plan-Apochromat 63x/1.4 NA oil-immersion objective.

The illumination source of widefield mode on CellDiscoverer 7 was LED (Colibri, Zeiss; wavelength for excitation light: 385, 470, 567 and 625 nm); t, the collected emission light of fluorophore was detected by a 4096 x 3008-pixel CMOS sensor (Axiocam 712, Zeiss). The illumination source of confocal mode (LSM-900) on Celldiscoverer 7 and LSM-700 was a four-channel-laser set (405, 488, 561 and 640 nm, Zeiss); the collected emission light after reflected beam splitter were detected by GaAsP-PMT detector. A summary of the imaging parameters used for each dataset can be found in Table S2.

### 2.12 Image processing and analysis

Fiji/ImageJ 1.46 software (NIH, USA) was used for image analysis. ROIs of hippocampal subregions were selected using NeuN or DAPI staining based on the Paxinos and Franklin Brain Atlas (Paxinos and Franklin, 2019).

For mean intensity and cell-counting analysis, images from both brain hemispheres per mouse were captured. In general, median filter (radius = 2) was applied through stacks to remove salt-and-pepper noise. Then maximum intensity projection (MIP) was performed to correct signal intensity variations caused by sample plane tilting in tile-scan.

Mean intensity could be measured in ROIs after this step. For cell counting, background subtraction (rolling ball method, radius = 50 pixels) was performed to enhance contrast. Then thresholds for each marker (calculated via Triangle method) were applied to create binary image. To reduce noise in binary images, further processing was applied in order of “Despeckle”, “Erode” x2, “Remove outliers, (radius = 2, threshold = 50)”, and “Dilate” x2. The target cells (e.g., microglia) were finally counted automatically as particles greater than 10 μm^2^ (Analyze > Analyze particles > 10-Infinity).

For microglia volume and synaptic pruning analysis, three Iba1-positive cells were imaged per brain region for each animal. In general, the median filter (radius = 2) was applied through stacks to remove salt-and-pepper noise. As shown in Figure 6, maximum intensity projection (MIP) and wand tool were applied to the Iba1^+^ channel to create outlines of selected microglial cells. ROI of the soma area was manually created with polygon-selection tool; the rest part of traced cell was recognized as ROI of microglial processes.

For 2D microglia morphological analysis, Sholl, skeletal and fractal analyses were performed on binarized microglia images (Figure S7, C-F). The detailed protocol has been described elsewhere (Strackeljan *et al*., 2025). In brief, for the Sholl analysis, the center of the cell body was selected with a point-tool; the radius of concentric circles was set to be from 4 μm to 60 μm with 2 μm intervals (Figure S7D). The number of times that the microglial branches intercepted each of the circles was calculated. For skeletal analysis, the binary microglia images were skeletonized (Figure S7E), then “analyze skeleton” function of ImageJ was applied to calculate branches, endpoints, averaged branch length and maximal branch length. For the fractal analysis, the same binary images were converted to outlines (Figure S7F). The FracLac plug-in was used to analyze the cells, with ‘box counting’ applied and the ‘grid design Num G’ set to 4. The fractal dimension (Db, a statistical measure of pattern complexity), lacunarity (a geometric measure of how a pattern fills space), circularity (how circular the microglial cell is), span ratio (longest length/longest width), and density (number of pixels/area) were measured.

Then, for 3D microglia volume and synaptic pruning analysis, the ROIs described above (Figure 6A) were applied to pre-smoothed images for stack-wise analysis. The threshold to recognize the Iba1^+^ area was first estimated with the Triangle method, then set as 20 to 255 for a binary mask to reduce the auto-threshold-variation among stacks and cells (Figure 6B). The volume of microglia was calculated as “cumulative area in ROIs through stacks” x “stack thickness”.

Regarding the synaptic pruning analysis, a similar protocol was applied to the same image set on CD68^+^ channel to characterize lysosomal structures within microglial somatic ROIs. The threshold to recognize CD68^+^ area was first estimated by the Triangle method, then set as 10 to 255 to create lysosomal masks in Z stacks (Figure 7A). The integrated density through lysosomal volume was used to evaluate the total amount of engulfed pre- and postsynaptic markers: ∑ (“area of CD68^+^ in a soma ROI at given Z section” x “mean intensity of marker at the same section” x “section thickness”).

### 2.13 Statistical analysis of IHC data

Statistical analysis of IHC measurements was performed with SigmaPlot software (Ver.13, Systat Software GmbH). Grubbs’ test was applied to determine outliers. If any outlier is detected, the outlier would be additionally confirmed by a manual review of the respective raw image. After outlier exclusion by the Grubbs’ test (P < 0.05), residual data were averaged for further statistical analysis. A normality test (Shapiro–Wilk method) and an equal variance test (Brown–Forsythe method) were applied to determine the appropriate statistical test. For data not passing the normality test and equal variation test, a transformation to either logarithmic scale or ranking was applied. Two-way ANOVA with the Holm–Šidák post hoc test was applied to evaluate effects of age and drug. Data and statistical results were plotted with GraphPad Prism software (Ver.8.0, GraphPad Inc.). In all statistical tests, a *P*-value < 0.05 was considered statistically significant. Detailed statistical information for each graph are indicated in the respective figure legend.

## 3. Results

### 3.1 NANA12 improves cognitive flexibility in the attentional set-shifting task

The ASST assesses cognitive flexibility by requiring mice to learn and adapt to changing reward contingencies (Figure 2A). As shown in Figure 2B, animals first learn a reward rule (compound discrimination), then undergo reversals and attentional shifts, evaluating their ability to renew learned associations.

To analyze ASST performance, we fitted the models described in the Materials and Methods section to the experimental data. The most informative results — treatment and age-related effects — are presented in Figures 2C-G. However, we also provide a more conventional Figure S2, which displays the estimated learning rates for each group. The distinction between effects and estimated values is crucial: effects represent differences between conditions, while estimated values reflect the total learning rate in a given group, obtained as the sum of effects. To aid interpretation, Figure S2 also includes *post-hoc* comparisons between groups, highlighting significant differences in learning rates and memory recall.

In the compound discrimination stage, young mice, regardless of NANA12 treatment, rapidly learned which medium was rewarded in direct trials, as evidenced by a strong learning rate in baseline control animals (log-odds coefficient for learning rate: 0.1, *P* < 10⁻⁶; Figure 2C). The absence of a significant effect of NANA12 in young mice indicates that their learning rate remained comparable to that of the control group (Figure 2C). Although no significant differences were observed in old mice, very old mice exhibited a substantially lower learning rate (effect size [ES]: -0.06 log-odds units, *P* < 0.01). Notably, this decline in learning performance was attenuated by NANA12 treatment (ES: 0.09, *P* < 0.05).

When the reward rule was reversed, young mice adapted rapidly (log-odds coefficient for learning rate: 0.18). While old mice did not show significant impairments in reversal learning, very old animals exhibited a pronounced impairment (ES: -0.11, *P* < 10⁻⁴). NANA12 treatment significantly ameliorated this deficit, increasing the learning rate during the reversal stage (ES: 0.11, *P* < 0.05). These results suggest that NANA12 treatment mitigates age-related learning deficits, enhancing both initial learning and cognitive flexibility in very old mice, bringing their performance closer to that of younger cohorts (Figure 2C). In Figure 2C, D, F, G, the reversal learning metrics are inverted relative to the values reported in Supplementary Tables S3, S4, and S5, meaning that more positive values indicate better reversal learning.

A similar trend was observed in the intradimensional shift stage, where the same stimulus dimension (medium) remained relevant, but novel exemplars were introduced (Figure 2D). Young mice maintained a high learning rate (log-odds: 0.12, *P* < 10⁻¹⁰) and an even stronger reversal learning rate (log-odds: 0.24, *P* < 10⁻¹⁹). As in the compound discrimination stage, old mice did not exhibit significant impairments compared to younger animals, while very old mice showed evident deficits in both direct learning (ES: -0.07, *P* < 0.05) and reversal learning (ES: -0.16, *P* < 10⁻⁵). NANA12 treatment significantly improved reversal learning performance in very old mice (ES: 0.09, *P* < 0.05; Figure 2D).

Following the intradimensional shifting, mice proceeded to the extradimensional shift stage. Unlike previous stages, where animals were assumed to have no intrinsic preference for any specific medium or odor at the start of the experiment, the extradimensional shift stage required consideration of priors regarding medium 3 and 4, which had been shaped during the intradimensional shift. Figure 2E presents the observed effects of the fitted model for these priors. Positive values indicate that at the onset of the extradimensional shift stage, mice expected M4 (media rewarded during the reversal stage of the intradimensional shift) to remain rewarded, whereas negative values reflect a preference for M3.

Model-derived coefficients and experimental data demonstrate that control young mice retained a strong memory for the rewarding status of M4 (log-odds: 0.98, *P* < 10⁻⁵). However, NANA12 treatment significantly reduced this memory recall (ES: -1.6, *P* < 10⁻⁶). Old mice exhibited weaker memory recall compared to control young mice (ES: -1.8, *P* < 10⁻³), while in the very old cohort, a similar but statistically non-significant reduction was observed. Notably, in both old and very old mice, NANA12 treatment significantly increased memory recall (ES: 2.9 and 2.6, respectively, *P* < 10⁻⁴ for both groups; Figure 2E).

At the beginning of the extradimensional shift stage, the medium dimension became irrelevant and was now rewarded at random. Despite this, mice initially held strong priors that M3 was not rewarded in the previous intradimensional shift reversal phase and exhibited learning behavior when it began to be occasionally reinforced in the direct phase of the extradimensional shift. This phenomenon is illustrated in Figure 2F.

As in previous phases, we analyzed the stimulus M3 that needed to be learned in the direct phase of the extradimensional shift, while M4, previously rewarded in the intradimensional shift reversal phase, was expected to be forgotten at this stage. The learning speed for medium 3-associated rewards in control young animals was positive (log-odds: 0.04, *P* < 10⁻⁵). Similar to memory recall, old mice exhibited a reduced learning speed in the direct stage compared to young controls (ES: -0.05, *P* < 0.05). However, NANA12 treatment significantly increased learning speed in both old and very old mice (ES: 0.12 and 0.14, respectively, *P* < 0.01 for both groups; Figure 2F).

The presence of a nonzero learning speed, approximately half that observed in previous stages, aligns with the fact that medium 3 was rewarded in only half of the trials due to its now-random reinforcement as an irrelevant dimension. This pattern suggests that a set-shift occurred during the direct learning stage, transitioning relevance from medium to odor. The absence of significant learning for medium reinforcement during the reversal stage (Figure 2F) further indicates that, at this stage, all experimental groups had successfully shifted their focus to odor as the new relevant dimension.

During the extradimensional shift stage, odor became the relevant dimension. Direct learning at this stage proceeded more slowly than in previous stages. Figure 2G presents the observed effects on odor learning speed. Modeling results showed that the learning rate for odor was 0.03 log-odds (*P* < 10⁻⁵) and was not significantly affected by age or treatment (Figure 2G). Thus, at this stage, mice were primarily engaged in “displacing” the previously relevant medium dimension, and group differences were most pronounced in this process (see Figure 2F). However, by the reversal stage, the medium dimension had lost significance for all experimental groups, leading to an increased learning speed for odors across all groups, regardless of age (log-odds: 0.06, *P* < 10⁻⁸). NANA12 treatment further enhanced this learning rate (ES: 0.06, *P* < 0.01; Figure 2G).

### 3.2. NANA12 improved performance of very old mice in Barnes maze

During the Barnes maze test (Figure 2A,B), mice were video-recorded and their trajectories were compared off-line between the groups, taking different parameters into consideration. One of key parameters characterizing the memory of escape hole location is the latency to reach the target zone during the probe trial (Figure 3C). Old mice showed good performance in this test and did not exhibit a significant difference from young mice. NANA12 did not change performance of young and old mice. However, very old mice from the control group exhibited higher latency compared to old control mice (*P* < 0.001). After treatment with NANA12, very old mice showed a great benefit from the treatment, displaying no significant difference from the young control group. Another notable finding was the total distance moved by the mice (Figure 3D). Old and young mice treated with either NANA12 showed no significant differences to controls in the total distance moved. However, very old control mice moved significantly less than young control and treated (*P* = 0.003) and old control (*P* = 0.02). In contrast, very old mice treated with NANA12 showed no difference compared to old and young mice. Analysis of mouse trajectories revealed that all NANA12-treated mouse groups preferably stayed near to the escape hole or two nearest holes during the probe trial (Figure 3E), in contrast to the very old control mice, which did not show clear preference.

These findings align with the results from ASST, further confirming that very old mice benefited the most from NANA12 treatment. The administration of NANA12 reduced the time required to reach the target zone and normalized the total distance moved, making it comparable to that of young and old mice.

### 3.3 NANA12 did not alter the size of hippocampal memory engrams in very old mice

To further explore the neural mechanisms underlying the cognitive improvements induced by NANA12, we quantified the spatial memory engram size defined here as the number of hippocampal c-Fos^+^ cells following memory retrieval during the Barnes maze probe trial.

As shown in representative images of c-Fos staining (Figure 4A) and by the quantitative analysis of c-Fos^+^ cells, there was a significant age-related reduction of c-Fos+ cells in CA1 *stratum pyramidale* (*P_age_* = 0.024, *F _(1,18)_* = 6.095), consistent with the known effects of aging on hippocampal function (Figure 4B). No significant effects of aging or NANA12 treatment were observed also in CA3 *stratum pyramidale* (*P_age_* = 0.196, *F _(1,18)_* = 1.806; *P_age x treatment_* = 0.980, *F _(1,18)_* = 0.0006) and in the dentate gyrus (DG) *stratum granulare* (*P_age_* = 0.771, *F _(1,18)_* = 0.0876; *P_age x treatment_* = 0.680, *F _(1,18)_* = 0.176). These results indicate that while aging significantly reduces c-Fos expression in CA1 *stratum pyramidale*, NANA12 treatment does not modulate the size of hippocampal memory engrams in very old mice. The cognitive benefits of NANA12 observed in behavioral tests are therefore more likely to be mediated by changes in the “quality” of hippocampal engrams or their usage downstream of the hippocampus rather than by changes in their size.

**Figure 4.**
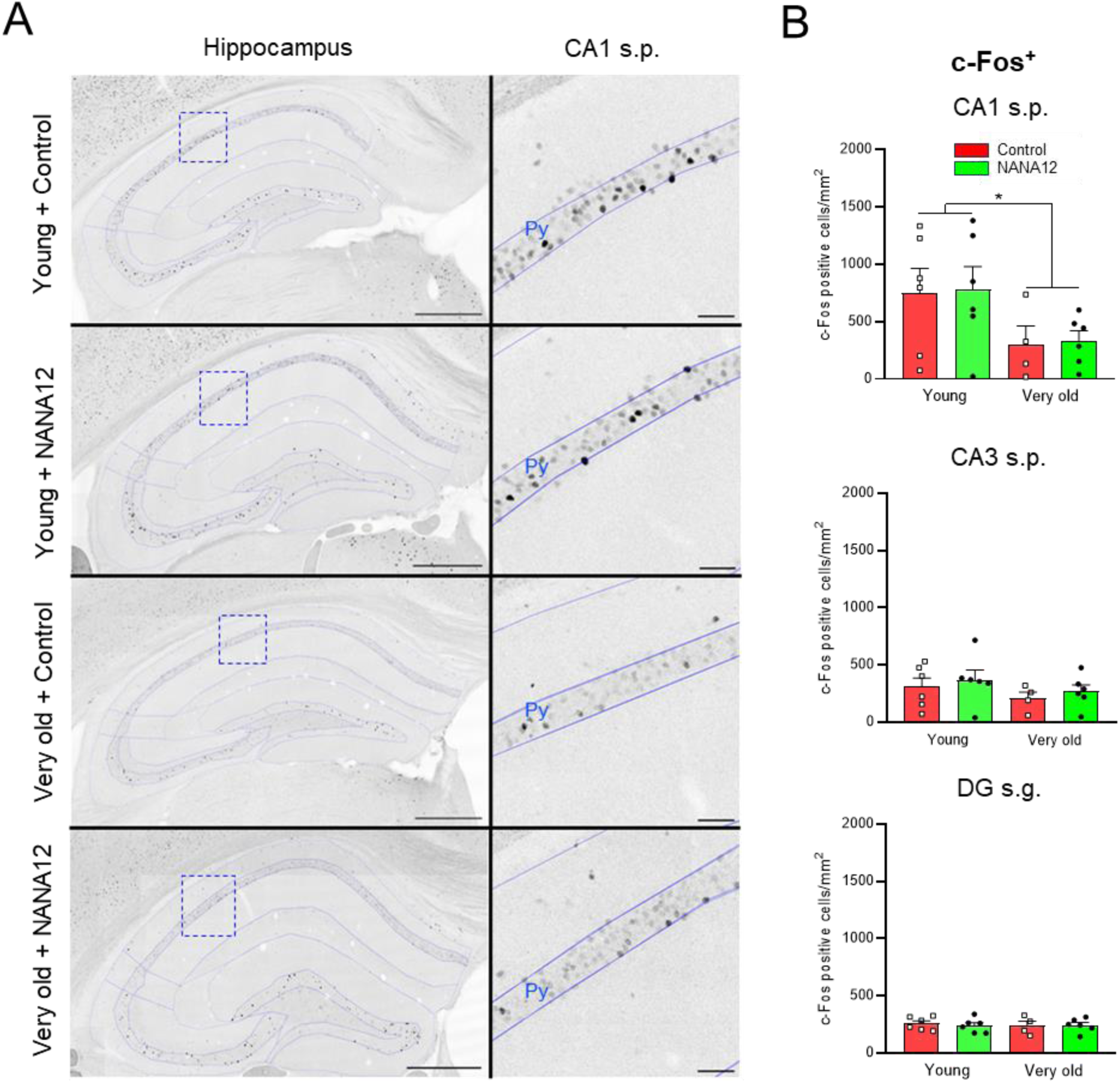
No effects of NANA12 on the size of c-Fos+ engrams in very old mice. (A) Representative images of c-Fos staining in different treatment groups. Left sub panels are tile-scanned panoramic images of hippocampal area. Scale bars: 500 μm. Right sub panels are high-magnification images in blue dotted frames in the left subpanels. Py (s.p.): pyramidal cell layer of the hippocampus. Scale bars, 50 μm. (B) Counts of c-Fos^+^ cells in CA1 s.p., CA3 s.p. and DG s.g. S.g.: granule cell layer of the dentate gyrus. Data are represented as mean ± SEM values. Two-way ANOVA, with Holm-Sidak post-hoc test; \**P* < 0.05.

### 3.4 No effects of NANA12 on dendritic integrity, TrkB signaling, polySia and lipofuscin expression

To assess whether NANA12 modulates dendritic integrity, we quantified MAP2 immunoreactivity across hippocampal subregions. As shown in Figure S3, no significant effects of aging or NANA12 treatment were observed in CA1 or DG subregion (*P* > 0.05 for all comparisons), indicating that dendritic structure remained stable across groups.

NANA12 has been reported to bind to BDNF (Kanato *et al*., 2008). To evaluate whether modulation of BDNF-TrkB signaling happens in a NANA12-dependent manner, we quantified p-TrkB immunoreactivity in the hilus where many p-TrkB+ cells have been reported (Jeanneteau *et al*., 2008). As shown in Figure S4, no significant effects of aging or NANA12 treatment were observed (P > 0.05 for all comparisons), suggesting that cognitive benefits attributed to NANA12 are not mediated by increased BDNF-TrkB signaling. To verify if NANA12 may affect the expression of endogenous polySia, thereby possibly competing for binding to polySia receptors, we performed polySia immunostaining and quantified its levels in hippocampal subregions (Figure S5). As previously reported, polySia staining was most prominent in mossy fiber projections in the CA3 *stratum lucidum* and hilus (Eckhardt *et al*., 2000). No significant effects of aging or NANA12 treatment were observed (P > 0.05 for all comparisons), indicating that NANA12’s cognitive benefits are unlikely to involve modulation of endogenous polySia expression.

Further, we quantified lipofuscin autofluorescence in various hippocampal subregions. Lipofuscin, a marker of cellular aging, accumulates in neurons over time and is thought to contribute to age-related functional decline. Representative images of lipofuscin auto-fluorescence revealed distinct patterns of accumulation across hippocampal layers, including the *stratum oriens*, pyramidal cell layer, *stratum radiatum* of CA1, and the granule cell layer, sub-granular zone, and the hilus (Figure 5A).

**Figure 5.**
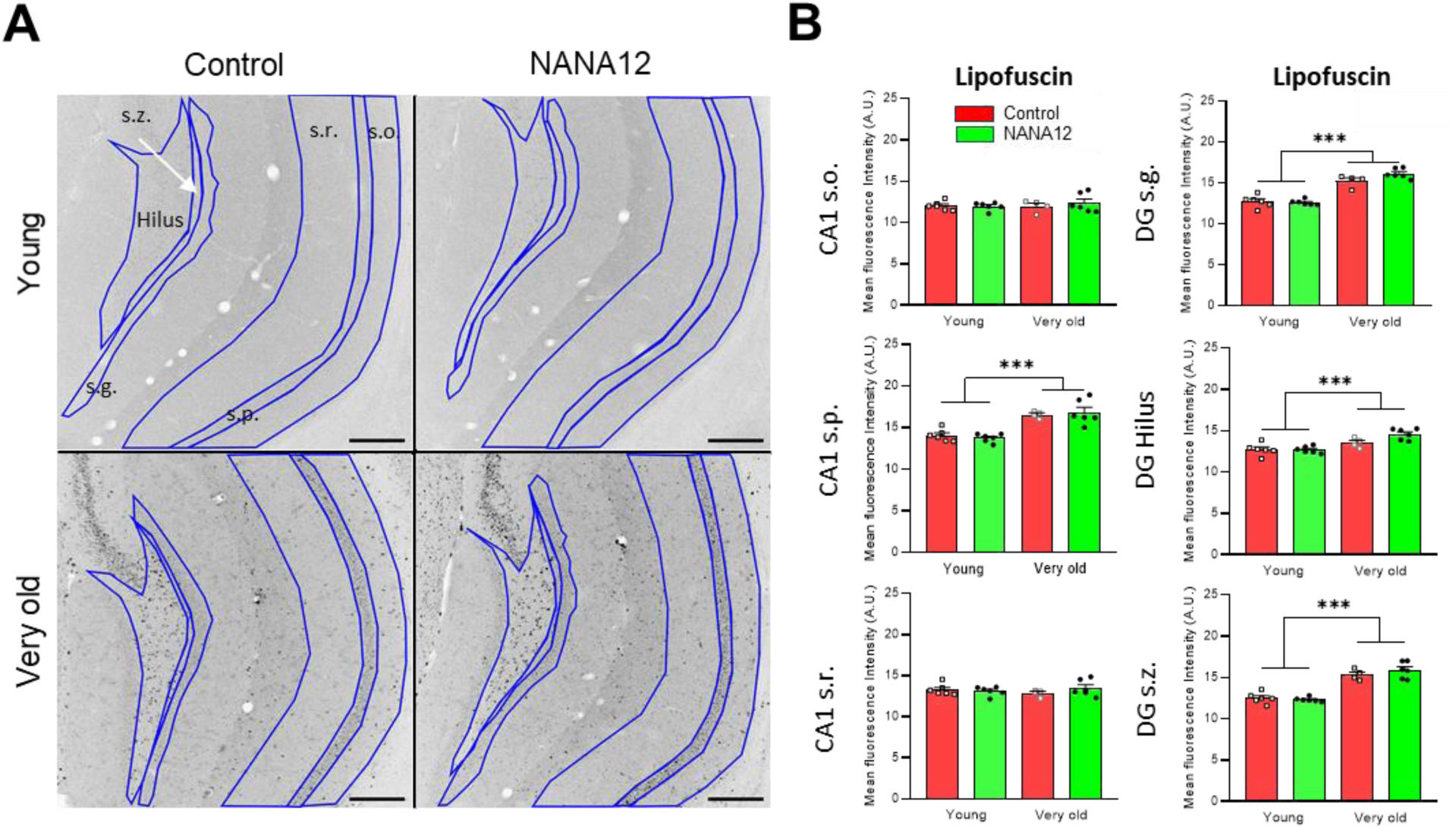
No effects of NANA12 on elevated lipofuscin levels in very old mice. (A) Representative images of Lipofuscin auto-fluorescence in different treatment groups. S.o.: oriens layer of the hippocampus; s.p.: pyramidal cell layer of the hippocampus; s.r.: radiatum layer of the hippocampus; s.g.: granule cell layer of the dentate gyrus; s.z.: sub-granular zone of the dentate gyrus. Scale bars: 200 μm. (B) Quantitation of Lipofuscin fluorescence intensity in different brain regions. Data are represented as mean ± SEM values. Two-way ANOVA, with Holm-Sidak post-hoc test; ***, *P* < 0.001.

Quantitative analysis of lipofuscin fluorescence intensity demonstrated a significant age-related increase in several hippocampal regions. In the CA1 *stratum pyramidale*, lipofuscin levels were markedly higher in very old mice compared to young mice (*P _age_* < 0.0001, *F_(1,18)_* = 48.539). Similarly, in the DG *stratum granulare* (*P _age_* < 0.0001, *F_(1,18)_* = 127.979), sub-granular zone (*P _age_* < 0.0001, *F_(1,18)_* = 110.237), and hilus (*P _age_* < 0.0001, *F_(1,18)_* = 25.823), lipofuscin signal was significantly elevated in very old mice. In contrast, lipofuscin levels in the CA1 *stratum oriens* (*P_age_* = 0.622, *F_(1,18)_* = 0.251) and *stratum radiatum* (*P_age_* = 0.901, *F_(1,18)_* = 0.016) showed no significant age-related changes. NANA12 treatment did not significantly reduce lipofuscin levels in any of the layers or subregions studied (*P_age x treatment_* > 0.05 for all regions). These results indicate that while lipofuscin accumulation is a prominent feature of aging in cellular layers in the hippocampus, short-term treatment with NANA12 does not mitigate this accumulation.

### 3.5 No major effects of NANA12 on the number and morphology of microglial cells

Microglia, the resident immune cells of the central nervous system, play a critical role in synaptic pruning and the maintenance of neural circuit integrity. Dysregulation of microglial function has been implicated in age-related cognitive decline, making these cell population a potential target for interventions aimed at enhancing brain health. To investigate NANA12 effects on microglial functions, we first assessed the impact of NANA12 on microglial cell density in the hippocampus. Representative images of Iba1 staining revealed a clear increase in microglia cell number with aging (Figure S6A). Quantitative analysis confirmed this trend, showing a significant age-related increase in Iba1-positive microglia density (*P _age_* = 0.0091, *F _(1,18)_* = 8.546). NANA12 treatment did not alter this trend, as microglia density in very old mice treated with NANA12 remained elevated (*P_age x treatment_* = 0.821, *F _(1,18)_* = 0.05) (Figure S6B). To further characterize microglia activation state, which is reflected by microglia morphology, we next performed a detailed 2D morphological analysis (Figure S7A-F) of microglia.

Skeletal and fractal analyses revealed no significant effects of aging or NANA12 treatment on branch complexity (*P* > 0.05 for all metrics; Figure S7I-Q). However, the Sholl analysis followed by three-way ANOVA uncovered a statistically significant effect of age (*P _age_* = 0.024, *F _(1,522)_* = 5.159), treatment (*P _treatment_* < 0.001, *F _(1,522)_* = 20.773) and interaction between age and treatment (*P _age x treatment_* = 0.013, *F _(1,522)_* = 6.204) on the number of branches as a function of distance from soma. Post-hoc analysis further confirmed the total number of branch intersections in young group was increased by NANA12 treatment (*P* < 0.001, *t* = 5.286). In the group of very old animals, there was a trend that NANA12 slightly increased intersections at a distance from soma center > 16 μm, but the change was not significant according to post-hoc test (*P* = 0.166, *t* = 1.387). Given the modest effect size of Sholl analysis and the limitations of 2D morphometrics in capturing spatial complexity, we next employed three-dimensional reconstruction to assess volumetric and topological features of microglia.

Figure 6A,B illustrates 3D volumetric analyses of microglial soma and arborizations in the CA1 region. Results revealed age-dependent morphological changes with layer-specific patterns. In the CA1 *stratum oriens*, microglial soma volume was significantly increased in very old control mice compared to young controls (*P_age_* = 0.036, *F _(1,18)_* = 5.148), suggesting activation of microglia. NANA12 did not significantly change the soma volume (*P_age x treatment_* = 0.169, *F _(1,18)_* = 2.057). A similar pattern was observed in *stratum pyramidale*, where aging markedly enlarged soma volume, while NANA12 treatment showed no effects (*P_age_* = 0.008, *F _(1,18)_* = 8.750; *P_age x treatment_* = 0.654, *F_(1,18)_* = 0.208). No age- or treatment-related changes were detected in *stratum radiatum* (*P_age_* > 0.05 for all comparisons). Notably, the volume of microglial arborizations remained unaltered across all layers and treatment groups (*P* > 0.05), suggesting that soma hypertrophy occurs independently of branch remodeling (Figure 6C).

**Figure 6.**
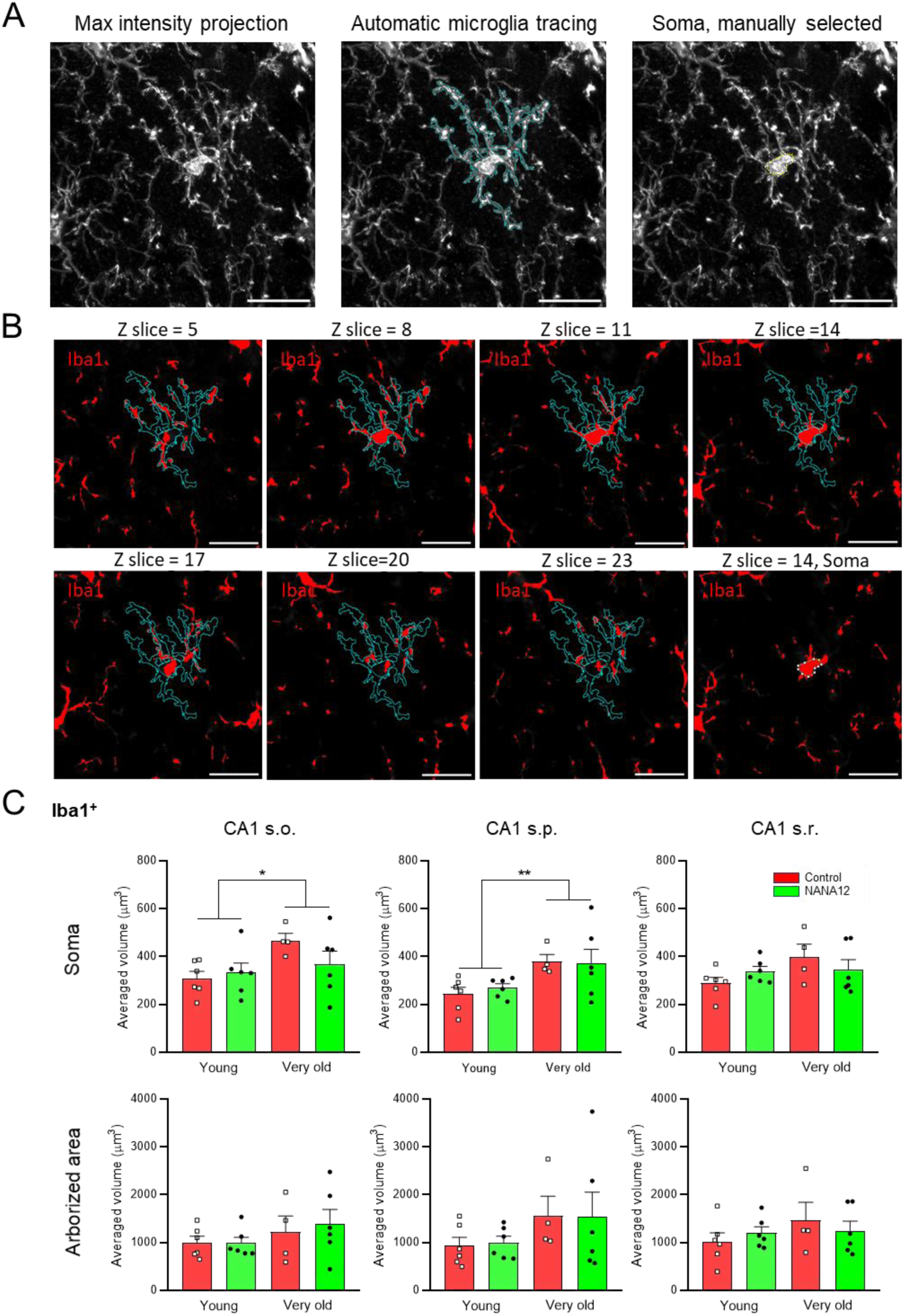
No effect of NANA12 on the size of microglia soma and arborizations. (A) Representative tracing of single microglial cell and its soma area on 2D projection plane. (B) Representative tracing of Iba1^+^ signal in 3D stack-wise manner. Scale bars: 20 μm. (C) Quantitation of averaged volume of single cell soma and arborizations in different layers of hippocampal CA1 region. 3 microglia cells from each region per animal have been randomly selected for averaging. Data are represented as mean ± SEM values. Two-way ANOVA, with Holm-Sidak post-hoc test; \**P* < 0.05; \*\**P* < 0.01.

### 3.7 NANA12 reduces lysosomal burden and synaptic phagocytosis by microglia in a layer-specific manner

To assess the impact of NANA12 on microglial lysosomal activity, we quantified the integrated fluorescence intensity of CD68, a marker of lysosomal compartments, within microglial soma across CA1 subregions (Figure 7A-B). In CA1 *stratum oriens*, aging markedly elevated CD68 intensity in very old control mice compared to young controls; NANA12 treatment significantly attenuated this increase in very old mice (Figure 7B (left); *P*_age_ = 0.0005, *F _(1,18)_* = 18.313; *P_age x treatment_* = 0.010, *F _(1,18)_* = 8.215). A similar age-dependent increase in CD68 intensity was observed in CA1 *stratum radiatum*. NANA12 treatment normalized CD68 levels in very old mice, suggesting a partial restoration of lysosomal homeostasis (Figure 7B (right), *P_age x treatment_* = 0.017, *F _(1,18)_* = 6.900). In contrast, NANA12 treatment could not reverse the age-related CD68 increase in *stratum pyramidale* (Figure 7B (middle), *P*_age_ < 0.0001, *F _(1,18)_* = 65.065; *P_age x treatment_* = 0.091, *F _(1,18)_* = 3.189). These analyses revealed significant age-related increases in lysosomal burden, with distinct spatial patterns of NANA12 therapeutic effects.

**Figure 7.**
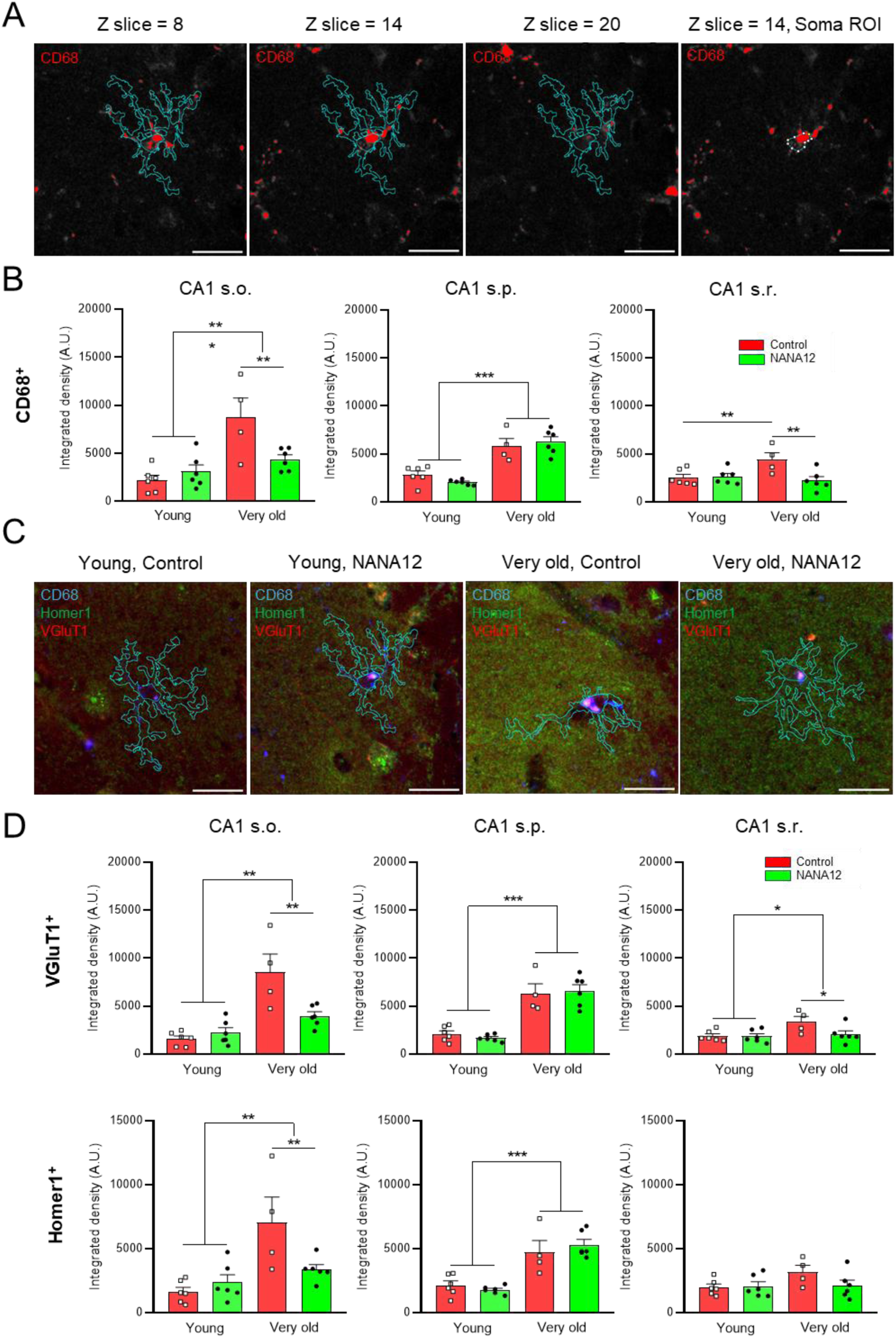
NANA12 reduced synaptic phagocytosis in the CA1 *str. oriens*. (A) Representative process to trace lysosomal CD68^+^ signal in 3D stack-wise manner corresponding to Figure 6A. Scale bars, 20 μm. (B) Quantitation of integrated fluorescence intensity of CD68 per microglial cell soma in different layers of hippocampal CA1 region. NANA12 reversed aging-induced CD68 increase in CA1 s.o. and s.r. regions but not in CA1 s.p. in very old mice. Data are represented as mean ± SEM values. Two-way ANOVA, with Holm-Sidak post-hoc test; \*\**P* < 0.01; \*\*\**P* < 0.001. (C) Representative single stack image showing synaptic markers (vGluT1 and Homer1) stained together with CD68 among all groups. Cell skeleton represents 2D projected single microglia ROI (Iba1^+^, as in Figure 6A). CD68^+^ ROIs extracted in soma area through Z-stacks were used for further quantitative analysis. Scale bars, 20 μm. (D) Quantitation of integrated fluorescence intensity of vGluT1^+^ and Homer1^+^ markers in CD68^+^ lysosomal structure in microglial soma in different sub-layers of CA1 region. NANA12 reduced integrated lysosomal vGluT1 and Homer1 intensity in the CA1 *stratum oriens* layer in very old mice. NANA12 reversed the increase on integrated lysosomal vGluT1 intensity but not for Homer1 in the CA1 *str. radiatum* layer in very old mice. In CA1 s.p., NANA12 did not reverse the age-related increase in vGluT1 and Homer1 in microglial somatic lysosomes in very old mice. 3 microglia cells from each region per animal have been randomly selected for averaging. Data are represented as mean ± SEM values. Two-way ANOVA, with Holm-Sidak post-hoc test; \**P* < 0.05; \*\**P* < 0.01; \*\*\**P* < 0.001.

To further investigate whether the effects of aging and NANA12 on microglial lysosomes are translated into changes in synaptic phagocytosis, we quantified the fluorescence intensity of synaptic markers (vGLUT1 and Homer1) within CD68_+_ compartments (Figure 7C-D). In CA1 *stratum oriens*, aging significantly increased lysosomal accumulation of both vGLUT1 (*P_age_* < 0.0001, *F _(1,18)_* = 32.332) and Homer1 (*P*_age_ = 0.0005, *F _(1,18)_* = 18.313). NANA12 treatment in very old mice reversed these increases, reducing both vGLUT1 intensity (*P_age x treatment_* = 0.023, *F _(1,18)_* = 6.145) and Homer1 (*P_age x treatment_* = 0.044, *F _(1,18)_* = 4.703). In CA1 *stratum radiatum*, age-related enrichment was observed for vGLUT1 (*P_age_* = 0.029, *F _(1,18)_* = 5.640) but not for Homer1 (*P_age_* = 0.126, *F _(1,18)_* = 2.569). NANA12 had a tendency to reduce vGLUT1 levels in very old mice in this layer (*P_age x treatment_* = 0.088, *F _(1,18)_* = 3.255), while the Homer1 intensity in very old mice remained unaffected by NANA12 (*P_age x treatment_* = 0.162, *F _(1,18)_* = 2.130). In CA1 *stratum pyramidale*, in contrast to other layers, NANA12 failed to attenuate age-related increases in lysosomal vGLUT1 (*P_age x treatment_* = 0.520, *F _(1,18)_* = 0.431) or Homer1 *(P*_age x treatment_ = 0.301, *F _(1,18)_* = 1.132) although both markers exhibited robust age-dependent accumulation (*P_age_* < 0.0001 for both vGLUT1 and Homer1). These findings demonstrate that NANA12 anti-phagocytic effects are layer-specific, preferentially targeting excitatory synapses in CA1 *strata oriens* and *radiatum*. Analysis of correlations between the individual behavioral performace in the Barnes maze and parameters characterizing the phagocytosis of synaptic markers vGLUT1 and Homer1 in the CA1 *stratum radiatum* revelead that mice performing worse (having longer latencies to the target area) had higher integrated intensities of vGLUT1 and Homer1 in CD68+ lysosomes (Figure S8) and this trend was present for very old control and NANA12-treated mice and even for young controls. These data support a vew that the beneficial effects of NANA12 on spatial memory are at least partially mediated by its modulation of synaptic phagocytosis.

## 4. Discussion

Our results show that treatment with NANA12 can improve cognitive flexibility and spatial memory recall abilities, particularly in very old mice. Furthermore, NANA12 may not be limited to enhance cognitive performance but also promotes locomotion and exploration of environment.

Considering humans, while some old individuals maintain stable cognitive function, others experience significant declines, particularly in cognitive flexibility. ASST studies indicate that older adults make more errors and show greater variability in reaction times than younger individuals, reflecting diminished executive control (Darna *et al*., 2024a; De Luca *et al*., 2003). Many individuals over 50 struggle with the extradimensional shift, a task requiring attention shifts to previously irrelevant dimensions, whereas younger adults typically complete this challenge with ease (Darna *et al*., 2024b). However, earlier ASST stages, such as intradimensional shifts, remain largely unaffected in aging (De Luca *et al*., 2003). Comparisons between middle-aged (40 years old) and elderly (70 years old) individuals confirm that aging selectively impairs extradimensional shifting while sparing simpler discrimination tasks (Owen *et al*., 1991). These findings suggest that deficits in cognitive flexibility primarily emerge in more complex set-shifting stages.

Despite well-documented cognitive decline in humans, mouse studies have not consistently shown similar impairments. No significant differences in ASST performance have been observed between young (∼5 months) and old (∼24 months) mice. Reports of increased response latencies during the extradimensional shift likely stem from non-cognitive factors, while performance was the same in both groups (Young *et al*., 2010). Similarly, old (23-month-old) and young (4-month-old) mice performed comparably across ASST stages, including the extradimensional shift (Tanaka *et al*., 2011). These findings suggest that traditional ASST analysis may lack the sensitivity to detect subtle age-related cognitive changes. Thus, while cognitive aging is evident in humans, ASST performance in old mice does not consistently reveal impairments, raising questions about the test’s ability to detect mild frontal cortex dysfunction in aging models.

To address these discrepancies, we adopted a more sensitive analytical approach, replacing trials-to-criterion analysis by trial-wise modeling and included very old mice (28 - 30 months). Traditionally, ASST data have been analyzed using trials-to-criterion, a widely accepted but potentially insensitive method. Recently, quasi-mechanistic reinforcement learning models have been introduced to enhance data interpretation (Talwar *et al*., 2024; Yearsley *et al*., 2021). These models explain behavior through reward prediction errors and learning rules, providing insight into choice adaptation over time. However, they pose challenges, including computational complexity, reliance on strong theoretical assumptions, and the need for large datasets. Additionally, reinforcement learning models often face identifiability issues, where different parameter settings yield similar behavioral outputs, complicating model comparison and validation. Their application to biological learning mechanisms remains an ongoing area of research.

Among available approaches, generalized linear mixed models (GLMMs) offer an optimal balance between computational efficiency and interpretability. Unlike trials-to-criterion, GLMMs retain trial-by-trial information, and unlike reinforcement learning models, they do not require extensive parameter tuning or strong assumptions about cognitive processes. By providing a statistically robust framework for evaluating cognitive flexibility, GLMMs ensure biologically meaningful findings. To our knowledge, this study is the first to apply GLMMs for trial-wise ASST analysis, offering a novel method suited to detect subtle cognitive impairments in aging. To validate our trial-wise GLMM approach, we directly compared trials-to-criterion and trial-wise modeling, assessing their sensitivity in detecting age-related effects as well as differences between reversal and direct learning (Supplementary Tables S3-S8).

Our results indicate that GLMM provides a more powerful and sensitive method for detecting trial-by-trial learning differences in the attentional set-shifting task and more refined analysis of learning rates, reversal learning, and interactions between age and NANA12 treatment, which were not captured by the trials-to-criterion analysis. The latter, while useful for summarizing learning performance, fails to capture subtle effects of age, reversal learning, and NANA12 treatment, making it less effective for detecting cognitive differences in aged mice. While previous studies suggested that ASST performance in old rodents does not mirror human cognitive decline (Tanaka *et al*., 2011; Young *et al*., 2010), our findings challenge this assumption by demonstrating clear impairments in old and very old mice when assessed with a more refined analytical approach. Our findings also demonstrate that NANA12 treatment enhances cognitive flexibility, particularly in old and very old mice, by improving learning and adaptation across different ASST stages.

Aging-related cognitive decline is thought to arise from disruptions in prefrontal cortex function, particularly in neural circuits governing attentional control and learning via a feedback. Consistent with this, our results indicate that very old mice struggle to adapt when a previously irrelevant dimension becomes relevant, mirroring deficits observed in elderly humans attempting extradimensional shifts (Darna *et al*., 2024a; Owen *et al*., 1991). This impairment likely reflects an age-related decline in the ability to suppress previously learned associations, a process reliant on prefrontal-striatal interactions (Floresco *et al*., 2009). Interestingly, NANA12 not only enhanced learning rates but also modulated memory recall: it reduced prior-based biases in young mice while strengthening memory retention in aged cohorts. This suggests that cognitive aging may not simply be a function of slower learning but also of altered memory stability, with NANA12 potentially positively fine-tuning this balance.

The fact that reversal learning in the extradimensional shift was consistently faster than direct learning across all age groups raises intriguing questions about age-related cognitive adaptation. While older mice were impaired in shifting to a new dimension, their ability to unlearn and reverse contingencies in the EDS phase remained relatively preserved. This aligns with reports that reversal learning may rely on distinct neural mechanisms compared to set-shifting.

One possible explanation is that reversal learning itself improves with experience, meaning that repeated exposure to reversals throughout the task may facilitate subsequent reversal performance. Another explanation is that a weaker memory of the rewarded stimulus in the direct extradimensional shift stage could, in principle, make reversal learning easier. However, while this factor might influence the number of trials to criterion, we consider it unlikely to have a substantial effect on learning speed, which we analyzed using a trial-wise GLMM approach. This distinction is important, as it highlights the advantage of our analytical method in isolating learning dynamics from the confounding effects of memory stability. NANA12 further accelerated reversal learning across all groups, suggesting a broader enhancement of cognitive flexibility rather than a selective rescue of age-related deficits.

The Barnes maze is a widely used task for assessing spatial learning and memory, heavily dependent on the hippocampal function. Aging-related impairments in Barnes maze performance have been widely reported, with older animals typically displaying increased latency to locate the escape hole, longer search paths, and more frequent errors (Barnes, 1979). The Barnes maze was originally designed for rats, and much of the research using this test focuses on this species. Only a few studies have examined age-related deficits in mice. Deficits in C57BL/6J mice have been reported at 12 months of age (Bach *et al*., 1999), and these impairments have been linked to declining hippocampal plasticity, synaptic loss, and impaired hippocampal-cortical connectivity (Rosenzweig and Barnes, 2003).

In our study, very old control mice (29 months) exhibited clear impairments in target zone navigation, as reflected by increased latency, consistent with previous findings on age-related hippocampal dysfunction (Burke and Barnes, 2006; Kennard *et al*., 2013). These impairments likely reflect age-related changes in navigation strategies, as older animals tend to rely more on egocentric, cue-based strategies rather than hippocampus-dependent allocentric strategies, which are typically more efficient in the Barnes maze (Lee *et al*., 2024). Human studies indicate a similar pattern, where older adults shift from hippocampal-dependent navigation to more striatal or prefrontal-mediated strategies, contributing to age-related declines in wayfinding abilities (Harris *et al*., 2012; Rodgers *et al*., 2012; Zhang *et al*., 2021). These shifts in the navigation strategy correlate with reduced hippocampal activation in fMRI studies, suggesting that rodents and humans share common neural mechanisms of age-related memory impairments (Antonova *et al*., 2009).

Importantly, these impairments were effectively rescued by NANA12 treatment, which restored both search efficiency and target preference in very old mice, bringing their performance in line with younger groups. NANA12-treated very old mice showed a preference for the correct target zone and took more direct paths to reach it, suggesting that they retained a more efficient allocentric navigation strategy.

Surprisingly, old control mice performed comparably to young mice, showing no significant impairments in spatial memory recall. This contrasts with studies reporting progressive deficits as early as 18–20 months (Barreto *et al*., 2010; Kennard and Woodruff-Pak, 2011). One possible explanation is that exposure to cognitive challenges before Barnes maze testing served as environmental enrichment, helping maintain spatial memory in old mice. Environmental enrichment is known to enhance cognitive function, promote hippocampal neurogenesis, and improve synaptic plasticity in aging rodents (Schmidt *et al*., 2022). Previous studies have shown that rats housed in enriched environments perform significantly better in the Barnes maze than standard-housed controls (Heimer-McGinn *et al*., 2020), and prior cognitive training can improve hippocampal plasticity and memory retention in aged animals (Kennard and Woodruff-Pak, 2011). Since all mice in our study underwent attentional set-shifting tasks before Barnes testing, this prior experience may have served as a form of enrichment, engaging compensatory mechanisms that helped old (23–24 months) mice to maintain performance.

A possible mechanism underlying beneficial cognitive effects of NANA12 involves the regulation of NMDA receptor activity. Endogenous polySia modulates extrasynaptic NMDA receptor activity, balancing synaptic and extrasynaptic signaling, which is crucial for synaptic plasticity and cognitive functions (Varbanov *et al*., 2023). By preventing excessive extrasynaptic NMDAR activation while reinforcing synaptic learning processes, NANA12 may enhance cognitive flexibility by optimizing NMDAR-dependent plasticity, particularly in prefrontal and hippocampal circuits. More broadly, our results indicate that aging disrupts not only cognitive flexibility but also the dynamic balance between memory persistence and adaptation. By facilitating set-shifting and reshaping memory biases — likely through its effects on synaptic plasticity — NANA12 emerges as a promising candidate for mitigating executive dysfunction in aging.

Immediate early genes such as c-Fos are widely used to assess neuronal activation and memory-related engram formation. While the c-Fos role in memory is well-established, the region-specific impact of aging on c-Fos+ engrams across hippocampal subregions (CA1, CA3, and DG) remains poorly understood, particularly during spatial memory and cognitive flexibility tasks. Our study provides some of the first evidence that aging selectively reduces c-Fos expression in CA1, while sparing CA3 and DG. This finding is noteworthy, as few published studies have reported global hippocampal c-Fos reductions with aging but have not systematically examined subregion-specific changes in the context of cognitive tasks. A potential explanation for this decline is age-related epigenetic repression of c-Fos transcription. A previous work (de Meireles *et al*., 2019) demonstrated that aging leads to histone modifications of the c-Fos promoter, including a decrease in H3K4me3 and H4K8ac, both markers of active transcription. These changes correlate with cognitive deficits, suggesting a loss of hippocampal plasticity. Interestingly, exercise reverses these modifications, restoring c-Fos transcription and improving memory performance (de Meireles *et al*., 2019).

Despite the age-related decline in CA1 c-Fos expression, NANA12 did not restore hippocampal engram size to youthful levels. Instead, its cognitive benefits suggest improved engram efficiency rather than increased neuronal activation. This is supported by our finding that NANA12 rescues cognitive deficits without altering hippocampal engram size. Given CA1’s role as a relay between the hippocampus and cortical regions involved in memory and decision-making (Yu and Frank, 2015), NANA12 may enhance hippocampal-cortical integration rather than locally increasing neuronal activation. This aligns with recent findings showing that functional dedifferentiation in the hippocampus contributes to age-related cognitive decline (Nordin *et al*., 2025). Specifically, a loss of distinct hippocampal connectivity gradients has been associated with impaired memory performance, suggesting that network-level disorganization is a key feature of cognitive aging.

Taken together, our findings identify CA1-specific reductions in c-Fos expression as an important and underexplored consequence of aging. While previous limited studies have documented global c-Fos decline with aging, our study is among the first to demonstrate a region-specific effect, particularly in CA1, and to suggest that cognitive benefits of NANA12 may arise from enhanced network integration rather than elevated local hippocampal recruitment. In other words, NANA12 cognitive benefits could stem from restoring functional network coordination rather than increasing c-Fos+ hippocampal engrams.

Immunohistochemical analysis revealed that NANA12 had no effects on dendritic integrity, expression of endogenous polySia and lipofuscin, and TrkB signaling. These are important data showing no toxic effects of NANA12 and indirectly supporting the view that NANA12 beneficial effects are mediated by inhibition of GluN2B-containing NMDARs rather than through upregulating expression of endogenous polySia or TrkB signaling.

In addition to glutamatergic dysregulation, aging as well as neurodegenerative diseases are associated with neuroinflammation and increased microglial activation, contributing to excessive synaptic pruning and cognitive impairment (Heuer *et al*., 2024; Hong *et al*., 2016; Li *et al*., 2023). Overactive microglia may engulf and phagocyte synapses, leading to reduced synaptic connectivity, particularly in the aging brain. This synaptic pruning mechanism is mediated by the complement system, which contributes to cognitive decline and impaired plasticity (Cangalaya *et al*., 2023; Shea and Villeda, 2024).

By characterizing microglia morphology, we were able to detect microgliosis and increased soma of aged microglia as hallmarks of microglia activation in very old mice, but no effects of NANA12 on these parameters were revealed. This is in line with previous studies suggesting that polySia with an average degree of polymerization higher than 12 ‒ NANA20 for human Siglec-11 (Shahraz *et al*., 2015) or NANA24 for murine Siglec-E (Thiesler *et al*., 2021) ‒ is necessary to efficiently modulate these microglial receptors.

A recent study indicates that extrasynaptic NMDARs promote synaptic remodeling through GluN2B-dependent release of postsynaptic/dendritic lysosomes (Grochowska and Kreutz, 2024) Thus, it is plausible to assume that an excessive activation of this pathway may boost extracellular proteolysis and increase synaptic tagging with complement proteins C1q and/or C3, triggering excessive synaptic phagocytosis. Also, excessive GluN2B signaling may interfere with the normal function of synapses and promote synaptic phagocytosis through multiple pathways such as calcium overload and calpain activation (Hardingham and Bading, 2010), nNOS signaling (Sattler *et al*., 1999), p38 mitogen-activated protein kinase (Soriano *et al*., 2008), and caspase activation (Li and Sheng, 2012). Taken together, our data suggest that targeting GluN2B-containing NMDARs and normalizing synaptic function by NANA12 reduces synaptic damage and phagocytosis and counteracts cognitive decline in aging and possibly in neurodegenerative conditions associated with aging.

## Funding

This work has been supported by grants from Bundesministerium für Bildung und Forschung (BMBF), project number 16LW0463K/Go-Bio-Initial CogniSia-2 to H.T. and 16LW0464/Go-Bio-Initial CogniSia-2 to A.D., and DFG (DI 702/10-1 to A.D.; CRC1436 projects A01 and A03 to M.F. and A.D.; GE 801/17-1 to RGS).

## Author contributions

A.A performed in vitro experiments and data analysis, T. F. performed data analysis, H.T. performed in vitro experiments, data analysis, supervised in vitro analysis, and wrote related sections of the manuscript, A.D. designed the study, supervised in vitro analysis, and wrote the manuscript.

## Declaration of Competing Interest

A.D., S.J., R.G.S., and H.T. have filed an international patent application on “Polysialic acid and derivatives thereof, pharmaceutical composition and method of producing polysialic acid”, WO2020025653A3. The remaining authors declare that the research was conducted in the absence of any commercial or financial relationships that could be construed as a potential conflict of interest.

## Acknowledgments

We thank Patricia Zarnovican for excellent technical support.

## Supplementary data

Supplementary tables and figures.

## Data availability

Data will be made available on request.

## Supplementary Tables

**Table S1.** Reagents used for immunohistochemistry.

| Antibodies | Dilution | Source | Catalog number | RRID |
| --- | --- | --- | --- | --- |
| Rat monoclonal anti-c-Fos antibody | 1:500 | Synaptic Systems | 226017 | AB_2864765 |
| Rabbit polyclonal anti-Iba1 antibody | 1:500 | FUJIFILM Wako Pure Chemical Corporation | 019-19741 | AB_839504 |
| Rat monoclonal anti-Mouse COMPOUND DISCRIMINATION68 antibody | 1:500 | Bio-Rad | MCA1957 | AB_322219 |
| Guinea pig polyclonal anti-Homer 1 antibody | 1:500 | Synaptic Systems | 160004 | AB_10549720 |
| Chicken polyclonal anti-VGLUT 1 antibody | 1:500 | Synaptic Systems | 135316 | AB_2619822 |
| Chicken polyclonal anti-MAP2 antibody | 1:500 | Millipore | AB5543 | AB_571049 |
| Rabbit polyclonal anti-phospho-TrkB (Tyr816) antibody | 1:200 | Millipore | ABN1381 | AB_2721199 |
| Guinea pig polyclonal anti-NeuN antibody | 1:500 | Synaptic Systems | 266004 | AB_2619988 |
| Mouse monoclonal anti-polySia antibody | 1:200 | Medizinische Hochschule Hannover | mab-735 | AB_2619682 |
| Goat anti-Rat IgG (H+L) Cross-Adsorbed Secondary Antibody, Alexa Fluor® 647 | 1:1000 | Thermo Fisher Scientific | A-21247 | AB_141778 |
| Goat anti-Rabbit IgG (H+L) Highly Cross-Adsorbed Secondary Antibody, Alexa Fluor® 546 | 1:1000 | Thermo Fisher Scientific | A-11035 | AB_2534093 |
| Goat Anti-Chicken IgY H&L preadsorbed, Alexa Fluor® 405 | 1:1000 | Abcam | ab175675 | AB_2810980 |
| Goat anti-Guinea Pig IgG (H+L) Highly Cross-Adsorbed Secondary Antibody, Alexa Fluor® 488 | 1:1000 | Thermo Fisher Scientific | A-11073 | AB_2534117 |
| Donkey Anti-Chicken IgY (IgG) (H+L) AffiniPure, DyLight® 405 | 1:200 | Jackson Immuno Research Labs | 703-475-155 | AB_2340373 |
| Goat anti-Chicken IgY (H+L) Secondary Antibody, Alexa Fluor® 488 | 1:1000 | Thermo Fisher Scientific | A-11039 | AB_2534096 |
| Goat anti-Mouse IgG (H+L) Highly Cross-Adsorbed Secondary Antibody, Alexa Fluor® Plus 405 | 1:500 | Thermo Fisher Scientific | A-48255 | AB_2890536 |
| **Dye for Nuclear stain** | **Dilution** | **Source** | **Catalog number** | **RRID** |
| DAPI (4',6-diamidino-2-phenylindole) | 1:400 | Thermo Fisher Scientific | D1306 | AB_2629482 |

**Table S2.** Parameters of imaging acquisition.

| **Immunostaining** | **Microscope** | **Imaging mode** | **Objective lens** | **Scan zoom** | **Pixel size x Z stack interval** |
| --- | --- | --- | --- | --- | --- |
| Rat x c-Fos, Gt x Rat AF647 | Celldiscoverer 7 | Widefield | Plan-Apochromat 20x/0.95 NA air. 0.5x Tube lens | 1.3 x | 0.229 x 0.229 μm |
| Lipofuscin autofluorescence | LSM-700 | Confocal | EC Plan-Neofluar 10x/0.3 NA air | 0.5 x | 0.625 x 0.625 x 10 μm |
| Rb x Iba1, Gt x Rb AF546 | LSM-700 | Confocal | Plan-Apochromat 63x/1.4 NA oil | 1.2 x | 0.082 x 0.082 x 0.7 μm |
| Rat x CD68, Gt x Rat AF647<br>Gp x Homer1, Gt x Gp AF488<br>Chk x VGLUT1, Gt x Chk AF405 | LSM-700 | Confocal | Plan-Apochromat 63x/1.4 NA oil | 1.2 x | 0.082 x 0.082 x 0.7 μm |
| Chk x MAP2, Dnk x Chk 405 | LSM-700 | Confocal | EC Plan-Neofluar 10x/0.3 NA air | 0.5 x | 0.625 x 0.625 x 10 μm |
| Rb x Iba1, Gt x Rb AF546 | LSM-700 | Confocal | EC Plan-Neofluar 10x/0.3 NA air | 0.5 x | 0.625 x 0.625 x 10 μm |
| Rb x pTrk-B (4°C, 72h),<br>Gt x Rb AF 546 (4°C, 24h) | Celldiscoverer 7 | Confocal | Plan-Apochromat 20x/0.95 NA air. 1x Tube lens | 1.0 x | 0.124 x 0.124 x 1.0 μm |
| Ms x PSA, Gt x Ms AF 405<br>NeuN | Celldiscoverer 7 | Confocal | Plan-Apochromat 20x/0.95 NA air. 0.5x Tube lens | 1.3 x | 0.225 x 0.225 μm |

**Table S3.** Analysis of compound discrimination using a generalized linear mixed model.

| Parameter | Explanation | Effect | SE | P-value |
| --- | --- | --- | --- | --- |
| directTrialCount:directMedLeft | learning rate | 0.09933 | 0.01919 | 2.55E-07 |
| reverseTrialCount:directMedLeft | reversal learning rate | -0.1832 | 0.02218 | 2.93E-16 |
| isOld:directTrialCount:directMedLeft | learning rate x isOld | 0.06393 | 0.03491 | 0.06725 |
| isDrug:directTrialCount:directMedLeft | learning rate x isDrug | -0.0212 | 0.03016 | 0.48192 |
| isVeryOld:directTrialCount:directMedLeft | learning rate x isVeryOld | -0.0634 | 0.02309 | 0.0061 |
| isOld:reverseTrialCount:directMedLeft | reversal learning rate x isOld | -0.0243 | 0.04074 | 0.55086 |
| isDrug:reverseTrialCount:directMedLeft | reversal learning rate x isDrug | -0.012 | 0.03214 | 0.70818 |
| isVeryOld:reverseTrialCount:directMedLeft | reversal learning rate x isVeryOld | 0.11661 | 0.02764 | 2.58E-05 |
| isOld:isDrug:directTrialCount:directMedLeft | learning rate x isOld x isDrug | -0.0349 | 0.04637 | 0.45234 |
| isDrug:isVeryOld:directTrialCount:directMedLeft | learning rate x isVeryOld x isDrug | 0.08561 | 0.03908 | 0.02861 |
| isOld:isDrug:reverseTrialCount:directMedLeft | reversal learning rate x isOld x isDrug | 0.09626 | 0.05095 | 0.05902 |
| isDrug:isVeryOld:reverseTrialCount:directMedLeft | reversal learning rate x isVeryOld x isDrug | -0.1087 | 0.04546 | 0.01685 |

**Table S4.** Analysis of intradimensional shift using a generalized linear mixed model.

| Parameter | Explanation | Effect | SE | P-value |
| --- | --- | --- | --- | --- |
| directTrialCount:directMedLeft | learning rate | 0.11545 | 0.01758 | 6.87E-11 |
| reverseTrialCount:directMedLeft | reversal learning rate | -0.2422 | 0.0262 | 7.09E-20 |
| isOld:directTrialCount:directMedLeft | learning rate x isOld | 0.03841 | 0.03478 | 0.269718 |
| isDrug:directTrialCount:directMedLeft | learning rate x isDrug | -0.0279 | 0.02426 | 0.250953 |
| isVeryOld:directTrialCount:directMedLeft | learning rate x isVeryOld | -0.0728 | 0.02873 | 0.011357 |
| isOld:reverseTrialCount:directMedLeft | reversal learning rate x isOld | 0.03886 | 0.04562 | 0.394454 |
| isDrug:reverseTrialCount:directMedLeft | reversal learning rate x isDrug | 0.05807 | 0.03328 | 0.081215 |
| isVeryOld:reverseTrialCount:directMedLeft | reversal learning rate x isVeryOld | 0.16059 | 0.0332 | 1.44E-06 |
| isOld:isDrug:directTrialCount:directMedLeft | learning rate x isOld x isDrug | -0.0354 | 0.04362 | 0.417746 |
| isDrug:isVeryOld:directTrialCount:directMedLeft | learning rate x isVeryOld x isDrug | 0.03688 | 0.03564 | 0.300963 |
| isOld:isDrug:reverseTrialCount:directMedLeft | reversal learning rate x isOld x isDrug | -0.0034 | 0.0564 | 0.951672 |
| isDrug:isVeryOld:reverseTrialCount:directMedLeft | reversal learning rate x isVeryOld x isDrug | -0.0951 | 0.04313 | 0.027646 |

**Table S5.** Analysis of extradimensional shift using a generalized linear mixed model.

| Parameter | Dimension | Explanation | Effect | SE | p-value |
| --- | --- | --- | --- | --- | --- |
| directMedLeft | Media | memory | -0.9841 | 0.2116 | 3.57E-06 |
| isOld:directMedLeft | Media | memory x isOld | 1.78677 | 0.49916 | 3.54E-04 |
| isDrug:directMedLeft | Media | memory x isDrug | 1.6084 | 0.32406 | 7.66E-07 |
| isSuper:directMedLeft | Media | memory x isSuper | 0.70009 | 0.46512 | 0.13247 |
| isOld:isDrug:directMedLeft | Media | memory x isOld x isDrug | -2.944 | 0.68915 | 2.05E-05 |
| isDrug:isVeryOld:directMedLeft | Media | memory x isVeryOld x isDrug | -2.5782 | 0.64563 | 6.80E-05 |
| directTrialCount:directMedLeft | Media | learning rate | 0.04071 | 0.009 | 6.55E-06 |
| reverseTrialCount:directMedLeft | Media | reversal learning rate | -0.0068 | 0.01071 | 0.52762 |
| isOld:directTrialCount:directMedLeft | Media | learning rate x isOld | -0.0586 | 0.02725 | 0.0316 |
| isDrug:directTrialCount:directMedLeft | Media | learning rate x isDrug | -0.0969 | 0.01946 | 7.00E-07 |
| isVeryOld:directTrialCount:directMedLeft | Media | learning rate x isVeryOld | -0.0308 | 0.02503 | 0.21906 |
| isOld:reverseTrialCount:directMedLeft | Media | reversal learning rate x isOld | -0.0412 | 0.03093 | 0.18264 |
| isDrug:reverseTrialCount:directMedLeft | Media | reversal learning rate x isDrug | -0.0092 | 0.02083 | 0.65948 |
| isVeryOld:reverseTrialCount:directMedLeft | Media | reversal learning rate x isVeryOld | -0.0031 | 0.01964 | 0.87501 |
| isOld:isDrug:directTrialCount:directMedLeft | Media | learning rate x isOld x isDrug | 0.12498 | 0.0395 | 0.00158 |
| isDrug:isVeryOld:directTrialCount:directMedLeft | Media | learning rate x isVeryOld x isDrug | 0.13627 | 0.03457 | 8.42E-05 |
| isOld:isDrug:reverseTrialCount:directMedLeft | Media | reversal learning rate x isOld x isDrug | -0.1315 | 0.19011 | 0.48928 |
| isDrug:isVeryOld:reverseTrialCount:directMedLeft | Media | reversal learning rate x isVeryOld x isDrug | 0.02575 | 0.04209 | 0.54068 |
| directTrialCount:directOdourLeft | Odor | learning rate | 0.0278 | 0.00622 | 8.26E-06 |
| reverseTrialCount:directOdourLeft | Odor | reversal learning rate | -0.0635 | 0.01094 | 7.79E-09 |
| isOld:directTrialCount:directOdourLeft | Odor | learning rate x isOld | 0.0015 | 0.01695 | 0.92927 |
| isDrug:directTrialCount:directOdourLeft | Odor | learning rate x isDrug | 0.01413 | 0.01307 | 0.27988 |
| isVeryOld:directTrialCount:directOdourLeft | Odor | learning rate x isVeryOld | 0.00286 | 0.01502 | 0.84904 |
| isOld:reverseTrialCount:directOdourLeft | Odor | reversal learning rate x isOld | -0.0232 | 0.03005 | 0.44091 |
| isDrug:reverseTrialCount:directOdourLeft | Odor | reversal learning rate x isDrug | -0.0627 | 0.02152 | 0.0036 |
| isVeryOld:reverseTrialCount:directOdourLeft | Odor | reversal learning rate x isVeryOld | 0.00438 | 0.01955 | 0.8229 |
| isOld:isDrug:directTrialCount:directOdourLeft | Odor | learning rate x isOld x isDrug | -0.0212 | 0.02463 | 0.38915 |
| isDrug:isVeryOld:directTrialCount:directOdourLeft | Odor | learning rate x isVeryOld x isDrug | -0.0211 | 0.0214 | 0.32312 |
| isOld:isDrug:reverseTrialCount:directOdourLeft | Odor | reversal learning rate x isOld x isDrug | -0.2436 | 0.18986 | 0.1996 |
| isDrug:isVeryOld:reverseTrialCount:directOdourLeft | Odor | reversal learning rate x isVeryOld x isDrug | 0.01772 | 0.04207 | 0.6737 |

**Table S6.** Trial-to-criterion (T2C) analysis of compound discrimination. To evaluate cognitive performance in the attentional set-shifting task, we compared two analytical approaches: the trial-to-criterion (T2C) method and a Generalized Linear Mixed Model (GLMM)-based trial-wise analysis. The T2C approach defines learning success as the first trial at which an animal maintains at least 80% accuracy over a 10-trial window. In contrast, the GLMM approach considers trial-by-trial learning dynamics, incorporating both fixed and random effects to model variability across individuals (see Materials and Methods). For T2C approach, we examined the effects of aging (*isOld*, *isVeryOld*), NANA12 treatment (*isDrug*), and reversal learning (*isRev*) on the trial-to-criterion values. The T2C method detected global effects in compound discrimination (p = 0.004) and intradimensional shift (p = 6 × 10⁻⁴), suggesting potential influences of age and reversal learning in earlier stages. However, in compound discrimination, only *isVeryOld* was significant (p = 0.015), with an effect size of 9.4, indicating that very old mice required significantly more trials to reach the criterion, consistent with age-related cognitive decline.

| Parameter | Effect | SE | P-value |
| --- | --- | --- | --- |
| Intercept | 15.8181818 | 2.10222277 | 10<sup>-9</sup> |
| isOld | 2.93181818 | 4.07093688 | 0.47388086 |
| isDrug | -3.73484848 | 2.91039563 | 0.20375077 |
| isRev | 3.38181818 | 3.04641021 | 0.27086639 |
| isVeryOld | 9.38181818 | 3.76057041 | 0.01503739 |
| isOld x isDrug | 4.98484848 | 5.50873293 | 0.36871371 |
| isOld x isRev | -0.13181818 | 5.79542822 | 0.98192013 |
| isDrug x isRev | 2.93484848 | 4.26531968 | 0.49374782 |
| isDrug x isVeryOld | -3.79848485 | 5.1278725 | 0.46139375 |
| isRev x isVeryOld | 0.16818182 | 5.58178744 | 0.97605127 |
| isOld x isDrug x isRev | 0.81515152 | 8.27200841 | 0.92179076 |
| isVeryOld x isDrug x isRev | -2.98484848 | 7.50152895 | 0.69195131 |

| Global F-test |  |  |
| --- | --- | --- |
|  | T2C | GLMM |
| **F** | $F_{(11,68)} = 2.87$ | $F_{(12,1680)} = 24.01$ |
| **P-value** | 0.004 | 10<sup>-49</sup> |

**Table S7.** Trial-to-criterion (T2C) analysis of the intradimensional shift. The T2C analysis detected global effects in the intradimensional shift (p = 6 × 10⁻⁴), but no specific predictors, including age and reversal learning, reached significance, despite the global effect. This suggests that while T2C captures overall variation, it lacks the sensitivity to isolate specific learning impairments.

| Parameter | Effect | SE | P-value |
| --- | --- | --- | --- |
| Intercept | 16 | 1.713293712 | $10^{-13}$ |
| isOld | 0.25 | 3.317779007 | 0.940156202 |
| isDrug | -1 | 2.371947739 | 0.674650844 |
| isRev | 2.545454545 | 2.422963204 | 0.297182571 |
| isVeryOld | 3.25 | 3.317779007 | 0.33077025 |
| isOld x isDrug | 5.35 | 4.489570584 | 0.237540718 |
| isOld x isRev | 0.704545455 | 4.692048069 | 0.88108499 |
| isDrug x isRev | -0.363636364 | 3.390705939 | 0.914910381 |
| isDrug x isVeryOld | 7.55 | 4.489570584 | 0.097219336 |
| isRev x isVeryOld | 3.871212121 | 4.970520678 | 0.438779261 |
| isOld x isDrug x isRev | -1.686363636 | 6.36844586 | 0.791964648 |
| isVeryOld x isDrug x isRev | -3.053030303 | 6.576310775 | 0.643953938 |

| Global F-test |  |  |
| --- | --- | --- |
|  | T2C | GLMM |
| **F** | $F_{(11,68)} = 3.54$ | $F_{(12,1640)} = 24.4$ |
| **P-value** | $6 \times 10^{-4}$ | $10^{-50}$ |

**Table S8.** Trial-to-criterion (T2C) analysis of the extradimensional shift. Although trial-to-criterion analysis (T2C) indicated a NANA12 effect in extradimensional shift, the absence of a significant global effect (p = 0.34) in the F-test limits the interpretability of this finding. This suggests that T2C lacks the sensitivity to capture learning dynamics in more complex cognitive shifts, where multiple cognitive processes are involved.

| Parameter | Effect | SE | P-value |
| --- | --- | --- | --- |
| Intercept | 25.72727273 | 2.281926761 | $10^{-13}$ |
| isOld | -9.227272727 | 5.817794541 | 0.120230842 |
| isDrug | -10.22727273 | 3.30682604 | 0.00351789 |
| isRev | -4.560606061 | 3.841055099 | 0.241768833 |
| isVeryOld | -3.727272727 | 4.418932171 | 0.403739514 |
| isOld x isDrug | 14.22727273 | 8.259187952 | 0.092318039 |
| isOld x isRev | 11.56060606 | 8.487213408 | 0.180420942 |
| isDrug x isRev | 7.838383838 | 5.18130833 | 0.137814616 |
| isDrug x isVeryOld | 10.22727273 | 7.341281485 | 0.17091411 |
| isRev x isVeryOld | 1.560606061 | 7.596908559 | 0.838231383 |
| isOld x isDrug x isRev | -21.33838384 | 11.89134687 | 0.079941878 |
| isVeryOld x isDrug x isRev | -5.338383838 | 11.27316985 | 0.638274758 |

| Global F-test |  |  |
| --- | --- | --- |
|  | T2C | GLMM |
| **F** | $F_{(11,42)} = 1.16$ | $F_{(12,1640)} = 5.74$ |
| **P-value** | 0.34 | $10^{-19}$ |

**Figure S1.**
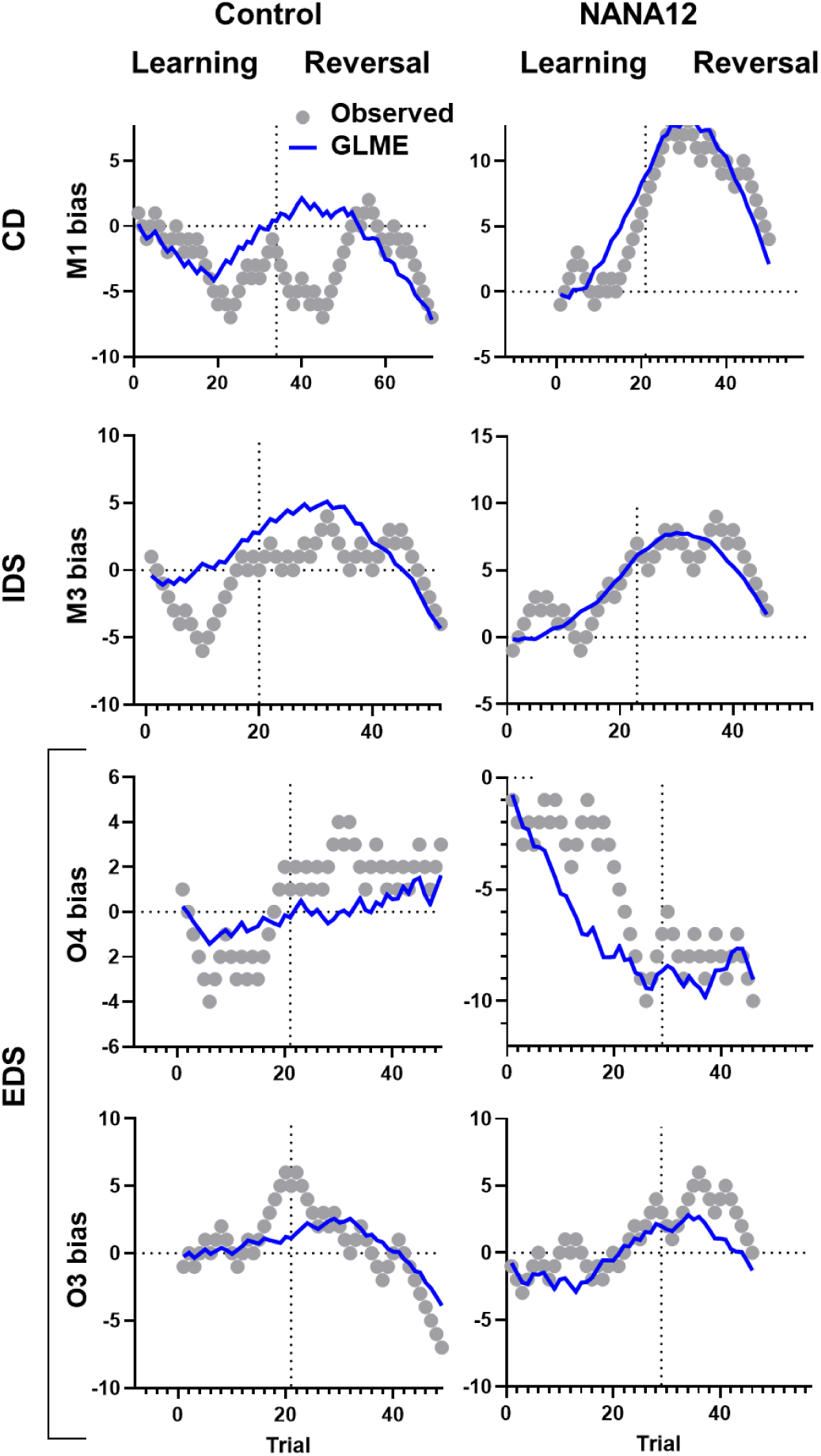
Representative stimuli bias trajectories for very old mice in control or after NANA12 treatment in each ASST stage. Related to Figure 2. The bias for each stimulus (y-axis) was calculated as the cumulative number of times the specified stimulus was chosen minus the cumulative number of times the opposing stimulus was chosen, up to each trial. Gray dots represent observed data, while the blue line indicates the GLME-predicted bias for the same mouse. Vertical dotted lines indicate the transition from the learning (direct) stage to the reversal stage. Horizontal dotted lines denote the bias = 0 reference. The y-axis scale is consistent within each row, while the x-axis scale is standardized across learning and reversal stages, resulting in “virtual” negative trial numbers in some rows. The figure shows representative data for very old control and NANA12-treated mice.

**Figure S2.**
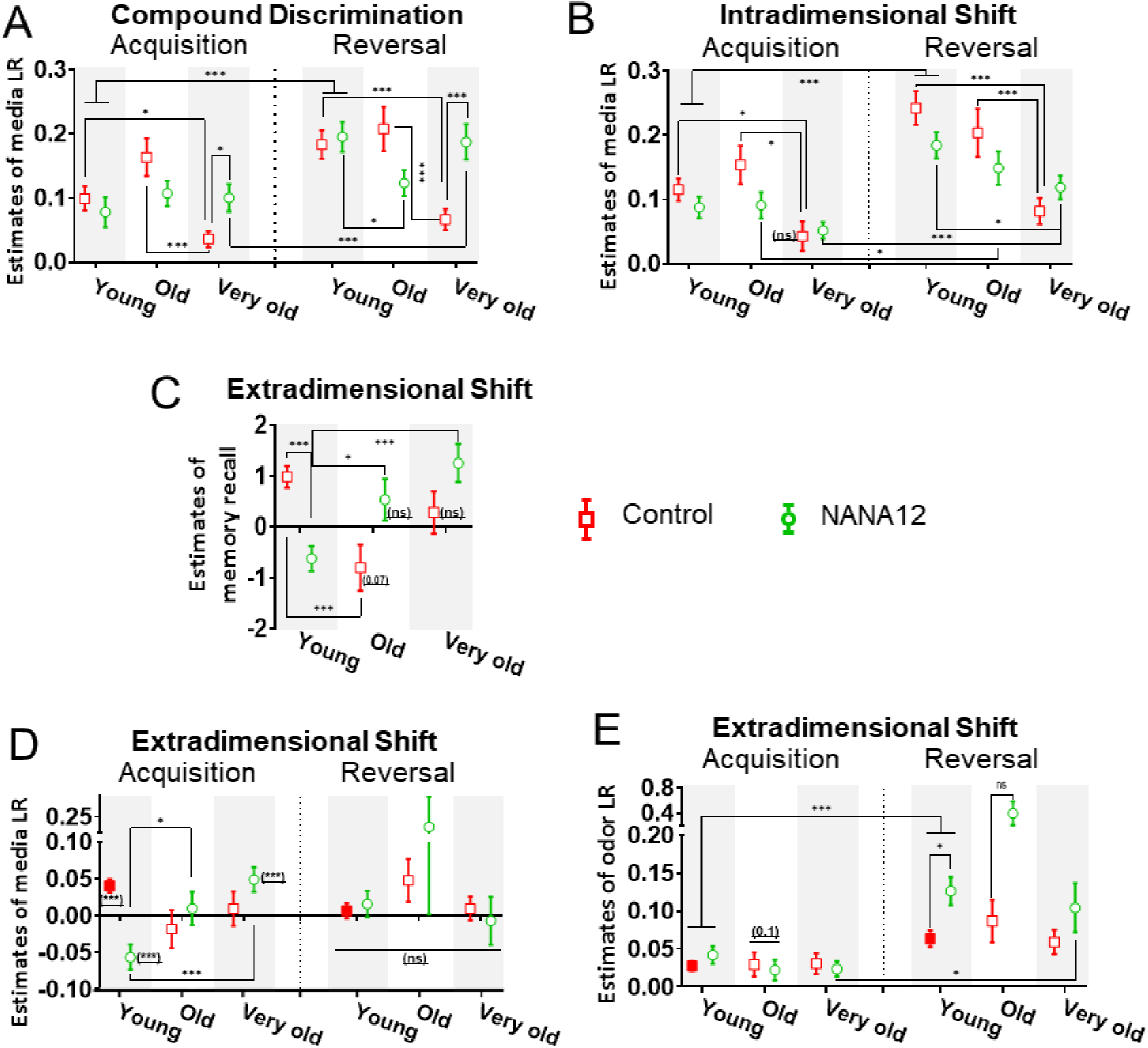
Estimated Learning Rates and Memory Recall Across ASST Stages. Related to Figure 2. To obtain direct estimates of learning rates and memory recall for each experimental group, rather than assessing the effects of age and NANA12 treatment, we reformulated our statistical models accordingly. For the compound discrimination **(C)** and intradimensional shift **(D)** stages, we modeled choice behavior as: isLeft ∼ 1 + ((directTrialCount + reverseTrialCount):directMedLeft):Age:isDrug + + *<u>(1+M1/3+O1/3|mouseID)</u>* For the extradimensional shift stage (E-G): isLeft∼-1 + ((directTrialCount + reverseTrialCount):directOdorLeft):Age:isDrug + + ((directTrialCount + reverseTrialCount):directMedLeft):Age:isDrug + + directMedLeft:Age:isDrug + *(1+M13+O13|mouseID),* Age and NANA12 were treated as categorical variables with full dummy variable coding. Multiple comparisons were corrected using the Benjamini-Hochberg false discovery rate correction. The models used to estimate learning rates and memory recall are structurally identical to those originally designed to assess the effects of age and NANA12 treatment. However, instead of treating age and treatment as primary factors of interest, we reformulated the models to express learning dynamics in terms of direct estimates for each group. These estimates include (C) learning and reversal learning rates for medium 1 and 2 in the compound discrimination stage, (D) learning rates for novel exemplars medium 3 and 4 in the intradimensional shift stage, (E) medium memory recall at the onset of the extradimensional shift, reflecting how strongly mice expected medium 4 (vs. 3) to be rewarded, (F) learning rates for media 3 and 4 in the extradimensional shift stage, and (G) learning and reversal learning rates for odors 3 and 4. Error bars indicate standard error. Significance thresholds are denoted as * *P* < 0.05, ** *P* < 0.01, ** *P* < 0.001.

**Figure S3.**
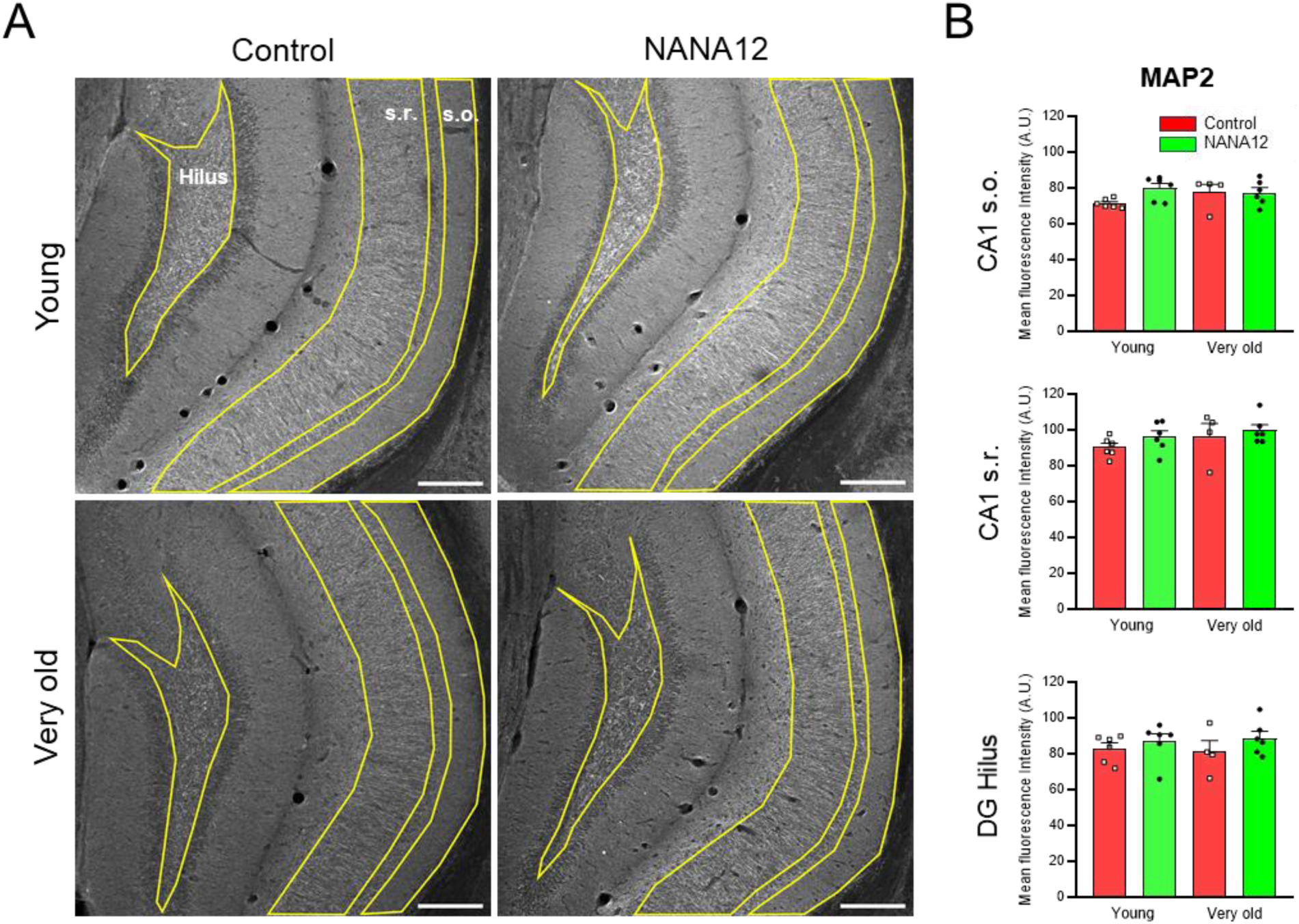
No effect of NANA12 on MAP2 immunoreactivity in the hippocampus. (A) Representative images of MAP2^+^ staining in all studied groups. ROIs were manually drawn according to the boarder of each sub-region. S.o.: oriens layer of the hippocampus; s.r.: radiatum layer of the hippocampus. Scale bars: 200 μm. (B) Quantitation of MAP2 fluorescence intensity in different brain regions. Data are represented as mean ± SEM values. No differences were detected by two-way ANOVA.

**Figure S4.**
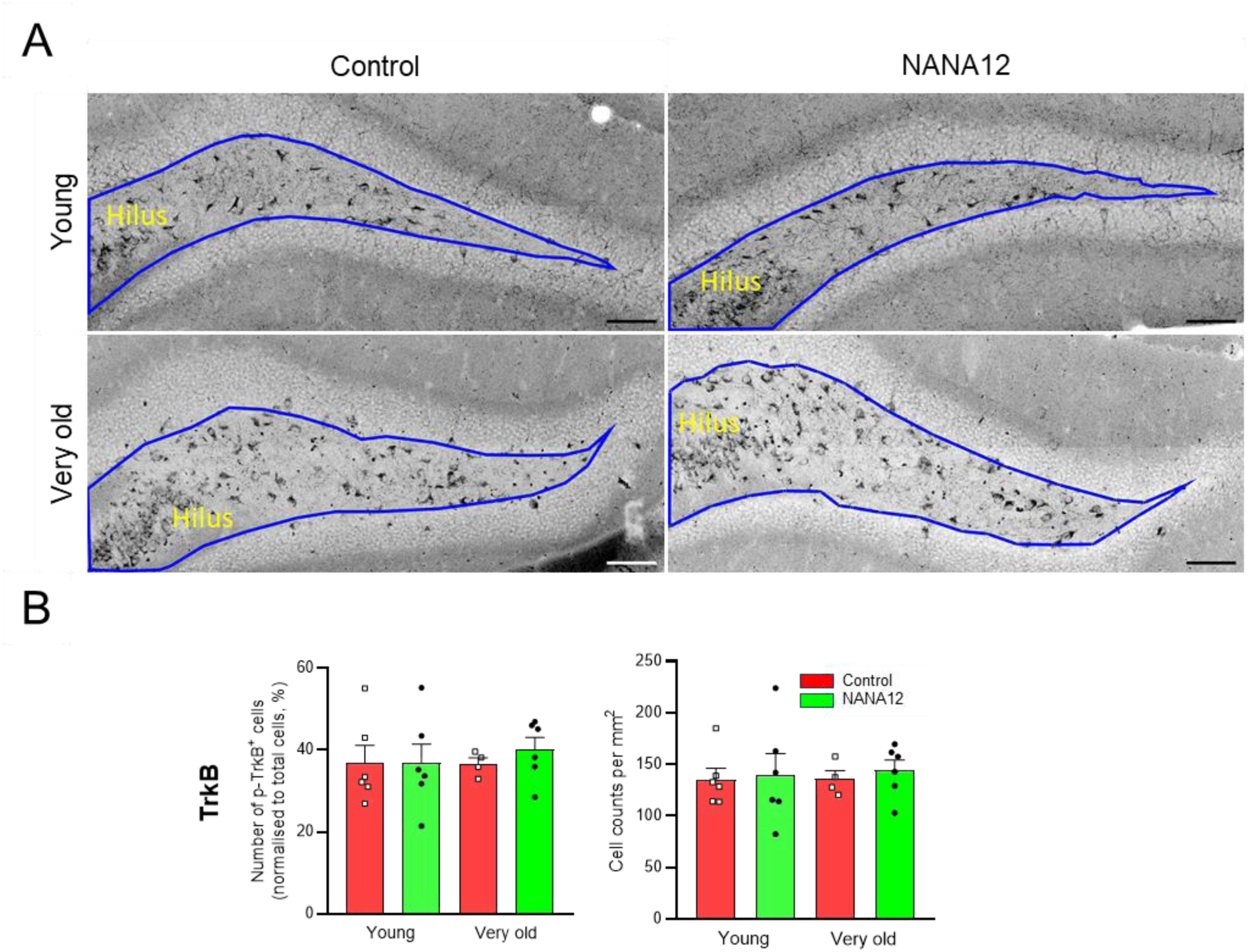
No effects of NANA12 treatment on the number of p-TrkB+ cells and mean intensity of p-TrkB in the hilus. (A) Representative images of p-TrkB^+^ staining in all studied groups. Hilus ROIs were manually drawn as in Figure S2. Scale bars: 200 μm. (B) Quantitation of p-TrkB^+^ fluorescence intensity in the hilus. Data are represented as mean ± SEM values. No significant changes were detected by two-way ANOVA.

**Figure S5.**
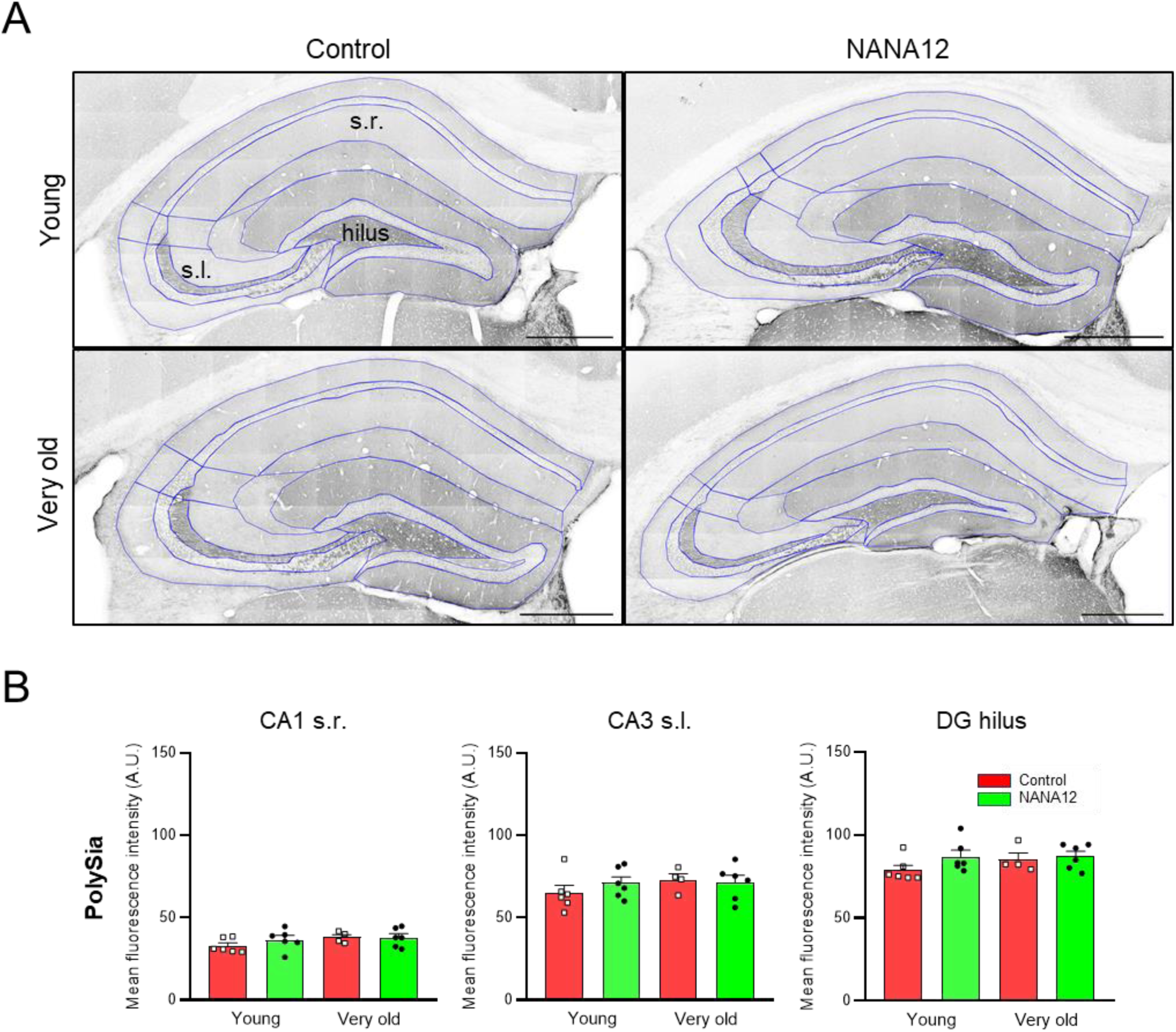
No effect of NANA12 on polySia immunoreactivity in the hippocampus. (A) Representative images of polySia^+^ stainings. Most of the ROIs were manually drawn based on NeuN staining, but ROI for CA3 s.l. was manually drawn according to the polySia staining. s.r.: *stratum radiatum*; s.l.: *stratum lucidum*. Scale bars: 500 μm. (B) Quantitation of polySia^+^ immunofluorescence intensity in CA1 s.r., CA3 s.l. and DG hilus. Data are represented as mean ± SEM values. No differences were detected by two-way ANOVA.

**Figure S6.**
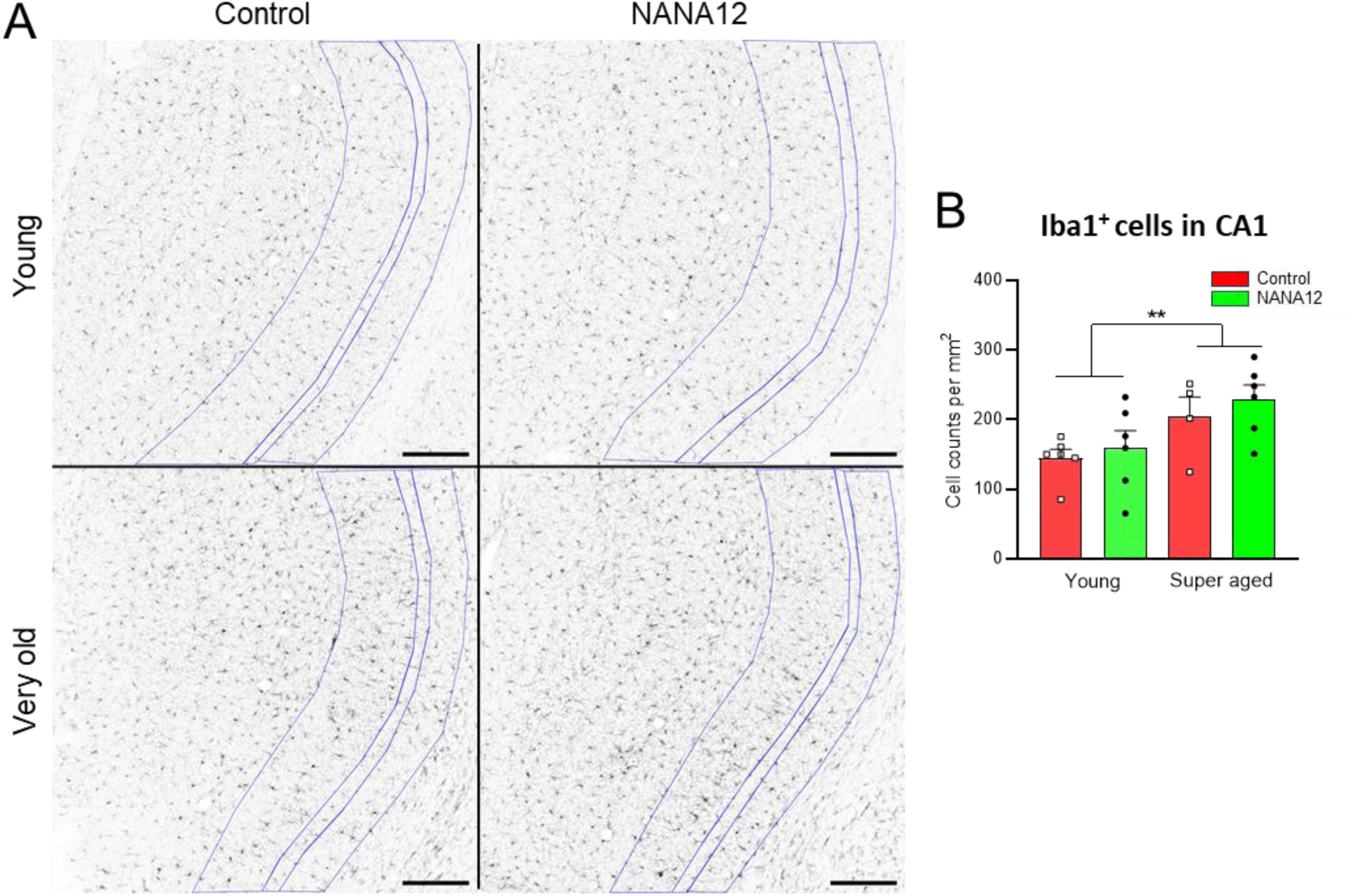
No effect of NANA12 on the number of microglial cells in the CA1 region. Related to Figures 6 and 7. (A) Representative images of Iba1^+^ staining in different treatment groups. ROIs of CA1 subregions were manually drawn according to co-stained Homer1 marker. Scale bars: 200 μm. (B) Number of Iba1^+^ microglia cells per mm^2^, cell number normalized to area of the selected CA1 region. Data is represented as mean ± SEM. Two-way ANOVA, with Holm-Sidak post-hoc test; **, *P* < 0.01.

**Figure S7.**
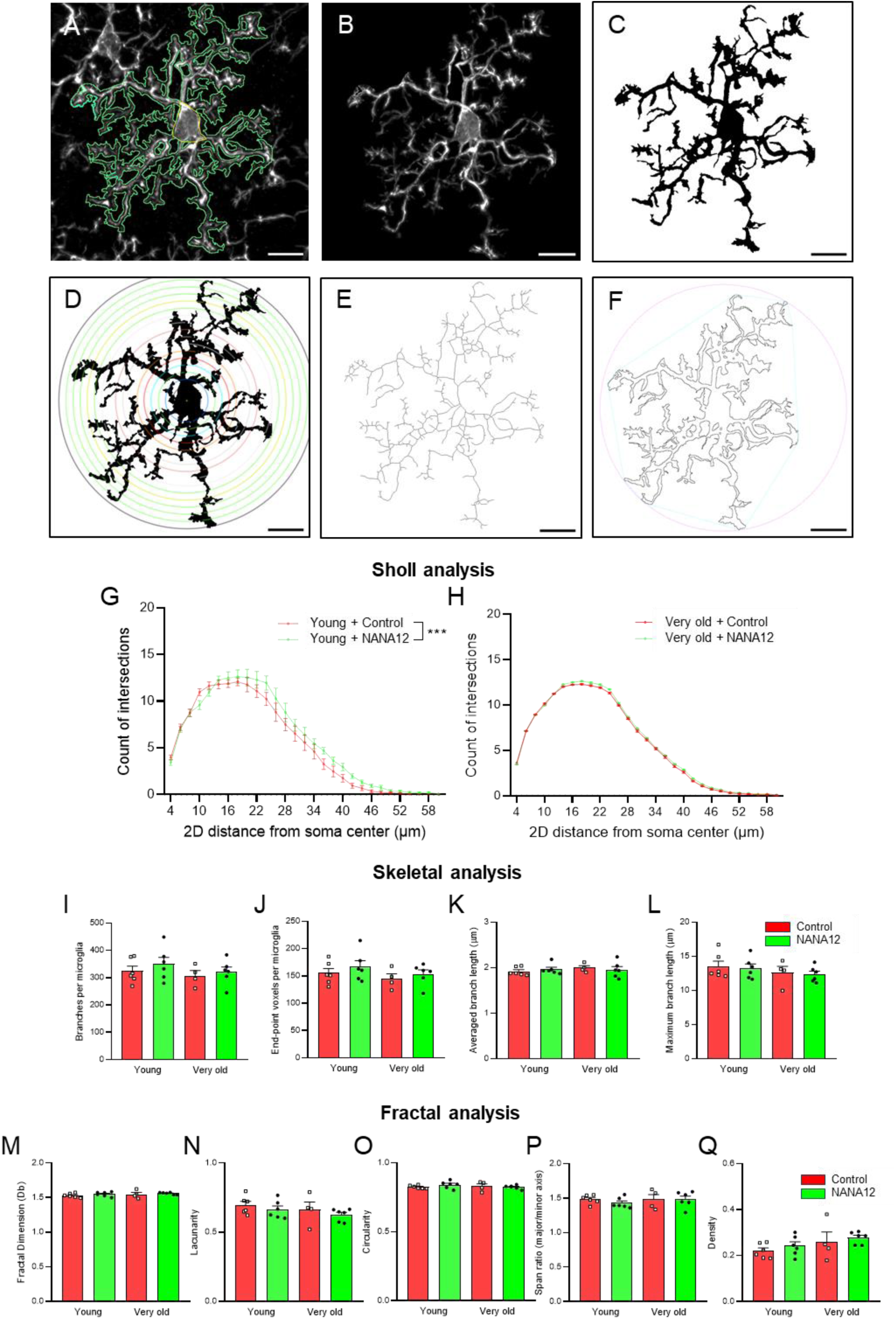
Minor effects of NANA12 treatment on microglial morphology in the CA1 region. Related to Figures 6 and 7. (A) to (F) Representative imaging to illustrate morphological analysis for single microglia. (A) Automatic tracing of Iba1^+^ microglia cell. (B) Selection of single microglia. (C) Binarized microglia image for further analysis. (D) Sholl analysis was performed to measure cell branching using concentric circles. Red circle indicates the circle with highest count of intersections with microglia. (E) For individual skeletal analysis, binarized microglia were skeletonized and analyzed using the “skeletal analysis” plugin. (F) Isolated microglia were converted to outlines and fractal analysis was performed using “FracLac” plugin. Scale bars: 10 μm. Results from Sholl analysis are presented in (G-H). Subpanels are (G) the distribution of intersections on concentric circles every 2 µm from the cell body for young group; and (H) the distribution of intersections on concentric circles for very old group. Results from skeletal analysis are presented in (I-L). The parameters are (I) number of branches; (J) number of branch endpoint pixels; (K) averaged branch length; (L) maximum branch length per microglia cell. Results from fractal analysis are presented in (M-Q). The parameters are (M) fractal dimension (Db) of microglia; (N) lacunarity of microglia; (O) circularity of microglia; (P) span ratio of microglia; (Q) pixel density per microglia. Results are presented as mean ± SEM. Two-way ANOVA, except for panel G and H (results are separately presented, two-way repeated measures ANOVA on ranks with Holm-Sidak post-hoc test; *, *P* < 0.05). Individual data points represent averaged results from 9 cells per mouse. In Sholl analysis, NANA12 presented a slight increase on distribution of intersections along distance to soma center in young mice and very old mice. *P _group_* = 0.0430, *F _(3,18)_* = 3.330; *P _group x distance_* = 0.029, *F _(84, 504)_* =1.348. No difference between groups was detected for all parameters acquired by Skeletal and Fractal analyses (*P* > 0.05).

**Figure S8.**
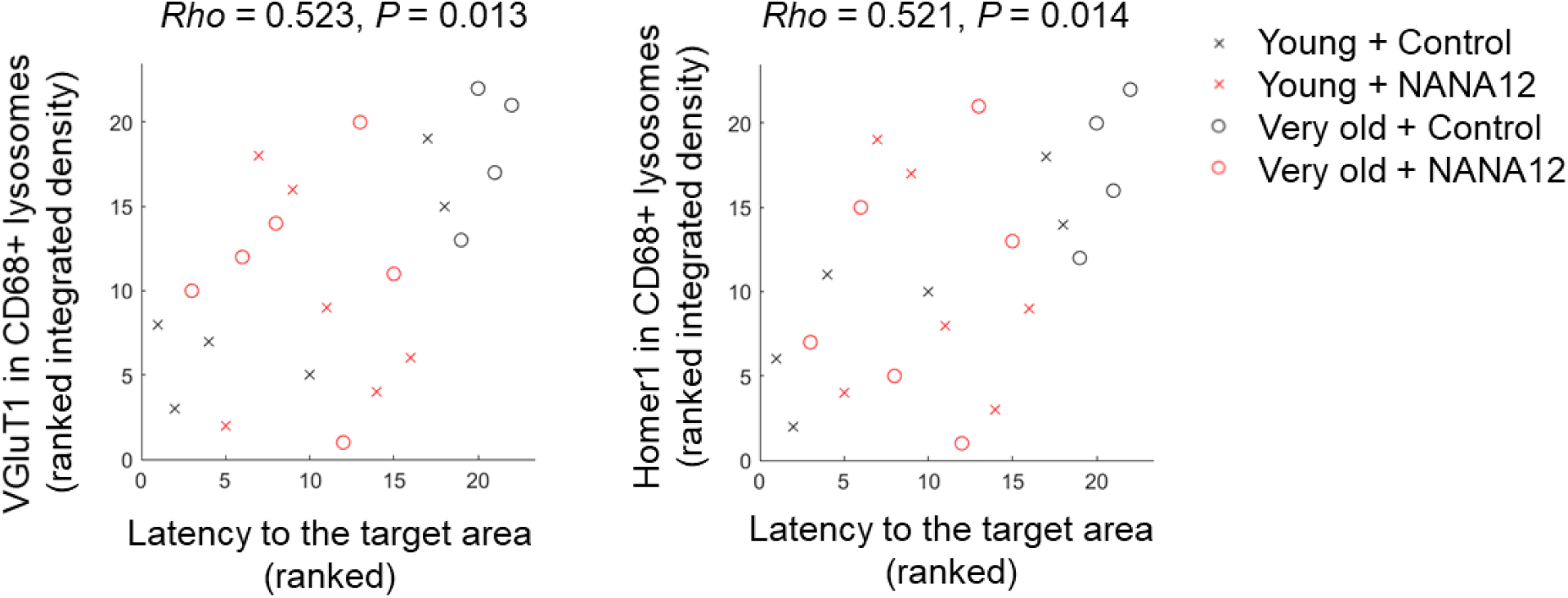
Correlation between the performance in the Barnes maze and synaptic phagocytosis. Related to Figure 7. Rho: Spearman coefficient of correlation.

